# A new take on model-based and model-free influences on mental effort and striatal prediction errors

**DOI:** 10.1101/2022.11.04.515162

**Authors:** Carolina Feher da Silva, Gaia Lombardi, Micah Edelson, Todd A. Hare

## Abstract

A standard assumption in neuroscience is that low-effort model-free learning is automatic and continuously employed, while more complex model-based strategies are only used when the rewards they generate are worth the additional effort. We present evidence refuting this assumption. First, we demonstrate flaws in previous reports of combined model-free and model-based reward prediction errors in the ventral striatum that likely led to spurious results. More appropriate analyses yield no evidence of a model-free prediction errors in this region. Second, we find that task instructions generating more correct model-based behaviour reduce rather than increase mental effort. This is inconsistent with cost-benefit arbitration between model-based and model-free strategies. Together, our data suggest that model-free learning may not be automatic. Instead, humans can reduce mental effort by using a model-based strategy alone rather than arbitrating between multiple strategies. Our results call for re-evaluation of the assumptions in influential theories of learning and decision-making.

## Introduction

A common view of decision-making posits that the human brain contains at least two distinct systems that guide behaviour: a model-free system, which repeats previous actions that frequently resulted in reward, and a model-based system, which plans to achieve rewarding outcomes based on a mental model of the environment [1]. The model-free system is proposed to operate automatically and support habit formation while the model-based system supports goal-directed behaviour. Model-based learning can lead to more rewards but is also more complex and thus is considered more effortful than model-free learning. The model-based strategy requires looking ahead and computing the possible outcomes of each action before making a decision, while the model-free strategy only requires retrieving averages of past outcomes from memory. The leading theory posits that decision-makers arbitrate between the two strategies according to a trade-off between reward and effort [2, 3]. If that trade-off is not favourable, the decision-maker will fall back on the model-free strategy. The model-free/model-based framework promotes precise computational definitions of behaviour that have been used to generate predictions for both healthy and psychiatric populations (e.g. [4, 5, 6, 7, 8, 9, 10, 11, 12, 13, 14, 15, 16, 17, 18, 3, 19, 20, 21, 22, 23, 24, 25, 26, 27, 28, 29, 30, 31]).

A large body of research investigates how model-based and model-free strategies interact using multi-stage decision tasks to dissociate the two strategies. The original two-stage task [4] is cleverly designed to determine whether an agent is using the correct model-based strategy or a simple model-free algorithm. In this task, agents make a choice at the first stage of each trial, then transition probabilistically to different second-stage states depending on their initial choice (Figure 1A). There are two options at the first stage as well as two second-stage states, and each first-stage option leads to one second-stage state with 70% probability (called a common transition) or to the other with 30% probability (called a rare transition). At the second stage, agents again make a choice between two options, and depending on their choice they may obtain a reward or not. The reward probabilities of the second-stage options change during the task (Figure 1C). This task can distinguish between model-free and model-based agents because the two types of agents have different probabilities of repeating their previous first-stage choice based on the previous trial’s events. Simple model-free agents are more likely to repeat a first-stage action that resulted in a reward regardless of the transition. In contrast, model-based agents use a model of the task’s probabilistic transition structure to make first-stage choices, being more likely to repeat rewarded actions after common transitions but less likely after rare transitions.

**Figure 1:**
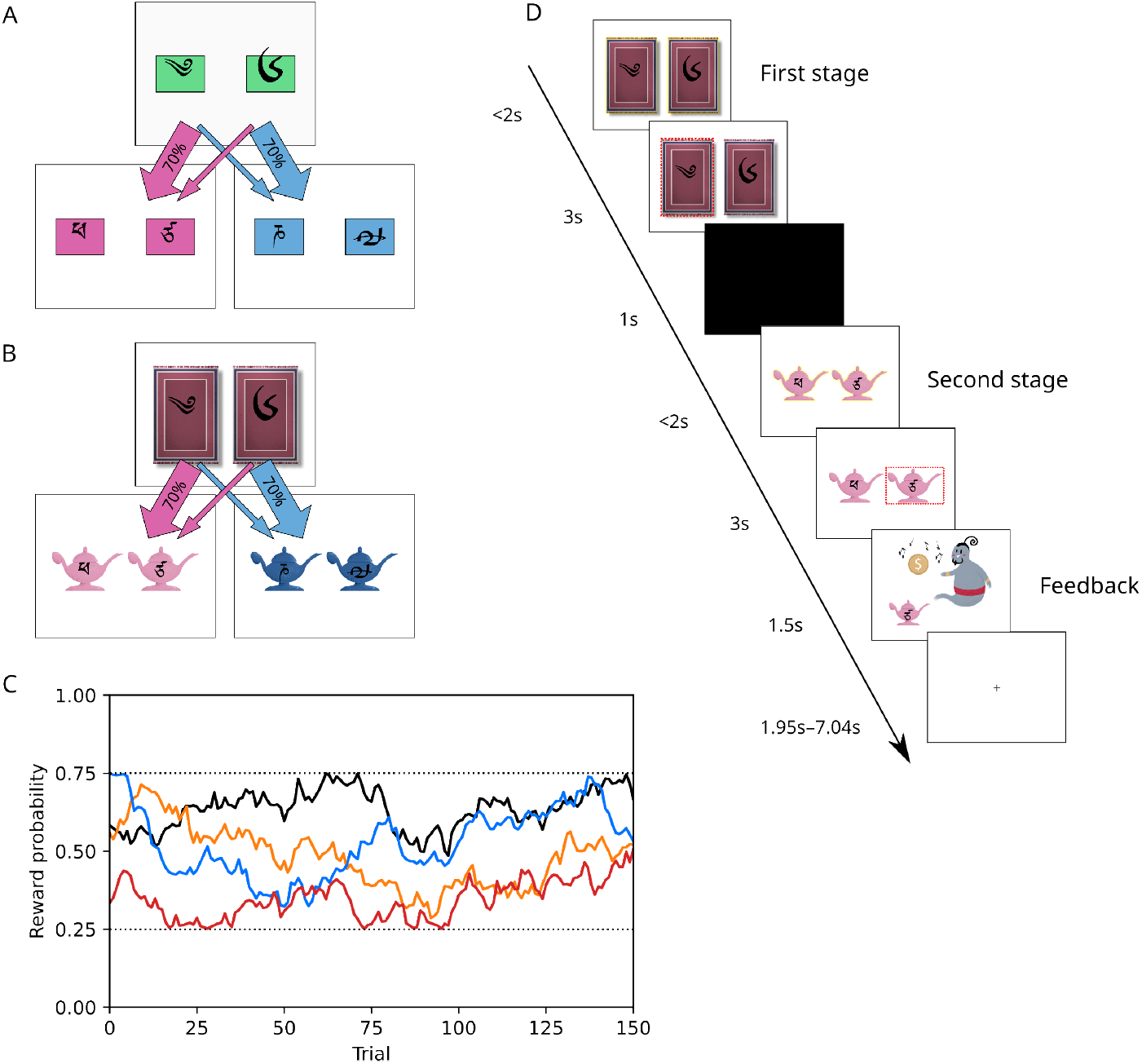
Structure and timeline of two-stage tasks. A) The original two-stage task [4] uses abstract symbols and common and rare transitions between states to distinguish between model-free and model-based strategies. In the first stage, participants choose one of two green symbols. Depending on which symbol they choose, they transition to the pink state or the blue state. In the second stage, participants choose one of two pink or blue symbols. B) The magic carpet task [28] has the same transition structure as the original two-stage task, but uses modified stimuli that allow for a detailed story in the instructions to improve understanding. The story instructions explained that the participant first chose a magic carpet and then flew on it to Pink or Blue Mountain. Both story and abstract instructions explained that when a lamp was chosen, a genie might or not come out and give the participant a coin. C) The reward probabilities of the four second-stage symbols change randomly between 0.25 and 0.75 during the task. D) Timeline of the two two-stage tasks used in this study, both using the same stimuli but either abstract or story-based instructions.

Past work has generally found that human behaviour in the two-stage task does not match either algorithm, instead seeming a hybrid mixture of the two [4, 5, 6, 7, 8, 9, 10, 11, 12, 13, 14, 15, 16, 17, 18, 3, 19, 20, 21, 23, 24, 25]. This apparent mixture matched functional magnetic resonance imaging (fMRI) results that seemed to show reward prediction errors (RPEs) resulting from both algorithms in the ventral striatum [4, 14, 26, 27, 31].

However, we demonstrated an exception to this apparently hybrid behaviour: when given detailed, story-based instructions, humans show nearly pure model-based behaviour [28]. This challenges the idea that behaviour in the two-stage task is the result of effort-based arbitration between simple model-free and correct model-based learning because the instructions don’t alter reward rates or the effort required by planning. This result is particularly surprising because the reward-effort trade-off in the original two-stage task favors model-free behaviour [3]. The hybrid hypothesis predicts that although behaviour might start off hybrid, it should shift towards model-free. However, studies have shown that strategies either do not change [25] or become more modelbased, not less, as humans practice the two-stage task over multiple days [13]. Furthermore, when rats were extensively trained to perform the two-stage task, they also developed a model-based strategy [22].

The instruction effect on behaviour suggests that humans did not actually use a hybrid strategy in previous studies. Instead, they may have misconceived how the two-stage task works and switched between incorrect strategies as their mental model of the two-stage task evolved during task performance. There is ample evidence that human participants mismodel the two-stage task [28, 23, 29] as well as other tasks with probabilistic transitions [32]. Critically, behavioural analyses of the two-stage task cannot distinguish between hybrid and incorrect model-based strategies [28], thus if participants form an incorrect model of the task, these analyses cannot identify the participants’ strategies.

Fortunately, the hybrid versus incorrect model-based hypotheses make additional predictions that can be used to test them. First, one can use neuroimaging to test which strategy best explains brain activity. The original paper [4] and several subsequent studies [14, 26, 27, 31] using the same two-stage task and analysis have reported evidence of combined model-free and model-based (i.e., hybrid) RPEs in the ventral striatum. Taken at face value, these results appear to support the hybrid hypothesis. However, we show that these results should not be taken at face value. First, the fMRI model used to produce these results computes an average of the RPE effects at the second stage and feedback, but without distinguishing between these two events one cannot show that model-free signals are encoded in the brain. Second, we mathematically prove that the apparent evidence for hybrid RPEs in the ventral striatum could be spurious. Critically, while we can replicate the previous results [4, 14, 26, 27, 31] in our data using the original model, if we remove the averaging there is no longer any indication of model-free or model-based RPEs in the ventral striatum. Third, one can also test how effortful participants found the task. The hybrid hypothesis predicts that people using a hybrid strategy exert less effort than those using a purely model-based strategy. Indeed, employing hybrid instead of model-based strategies reduces computational costs when robots navigate [33]. Therefore, if participants given the story instructions use a modelbased strategy, and those given the standard instructions use a hybrid strategy, then the story group will exert more mental effort. In contrast, the incorrect models hypothesis predicts that the story instructions help participants form and maintain the correct task model, while the standard instructions lead participants to spend more effort forming and arbitrating between inaccurate models. Thus, the two hypotheses make opposite predictions about effort.

We tested these predictions about effort and RPEs by randomly assigning healthy adult participants to different instruction conditions and quantifying their behaviour, pupil diameter, and brain activity during the task. The instruction conditions were (1) story, in which participants received elaborate instructions as a detailed story [28], and (2) abstract, in which participants received simple, abstract instructions as in past studies [4]. Our results show that participants in the abstract condition found the task more effortful and their understanding was lower compared to the story condition. When receiving feedback, the abstract group also exhibited larger pupil diameter, an effect previously linked to mental effort and uncertainty, and higher BOLD activity in prefrontal areas previously linked to strategy arbitration. Moreover, contrary to the hybrid hypothesis, simple model-free RPEs did not match activity patterns in ventral striatum. Overall, our findings suggest that participants in the abstract condition were not using a less effortful mixture of correct model-based and simple model-free learning. Instead, they were searching through various strategies and figuring out how the task works, which was more effortful than the approximately correct model-based strategy maintained by the story group.

## Results

Participants made decisions in an fMRI scanner in one of two conditions: abstract (*N* = 48) or story (*N* = 46; see Figure 1). We tested if the instructions changed behaviour, understanding, effort, and physiological responses during the task.

### Behavioural results

#### Story instructions lead to more correct model-based behaviour

We first tested if we could replicate previous findings that participants in the story condition behaved in a more correct model-based way compared to the abstract condition [28]. We analyzed decisions using the two standard approaches, 1) logistic regression on the stay probability in consecutive trial pairs, and 2) a hybrid model-based/model-free reinforcement learning (RL) model [4]. The logistic regression model describes the probability that a participant will stay with the same first-stage choice in a pair of consecutive trials as a function of the first trial’s outcome (reward or no reward), transition (common or rare), and their interaction [4]. Simple model-free agents show only a positive reward effect, correct model-based agents show only a positive reward-by-transition interaction, and hybrid agents show both. The results (Figure 2) indicate that in the story compared to abstract condition, the reward coefficient was smaller (0.999 posterior probability; mean difference: −0.21, 95% CI [−0.34, −0.08]) and the reward-by-transition interaction coefficient was larger (0.995 posterior probability; mean difference: 0.41, 95% CI [0.10, 0.72]). This suggests that the story instructions led participants to use a more correct model-based strategy.

**Figure 2:**
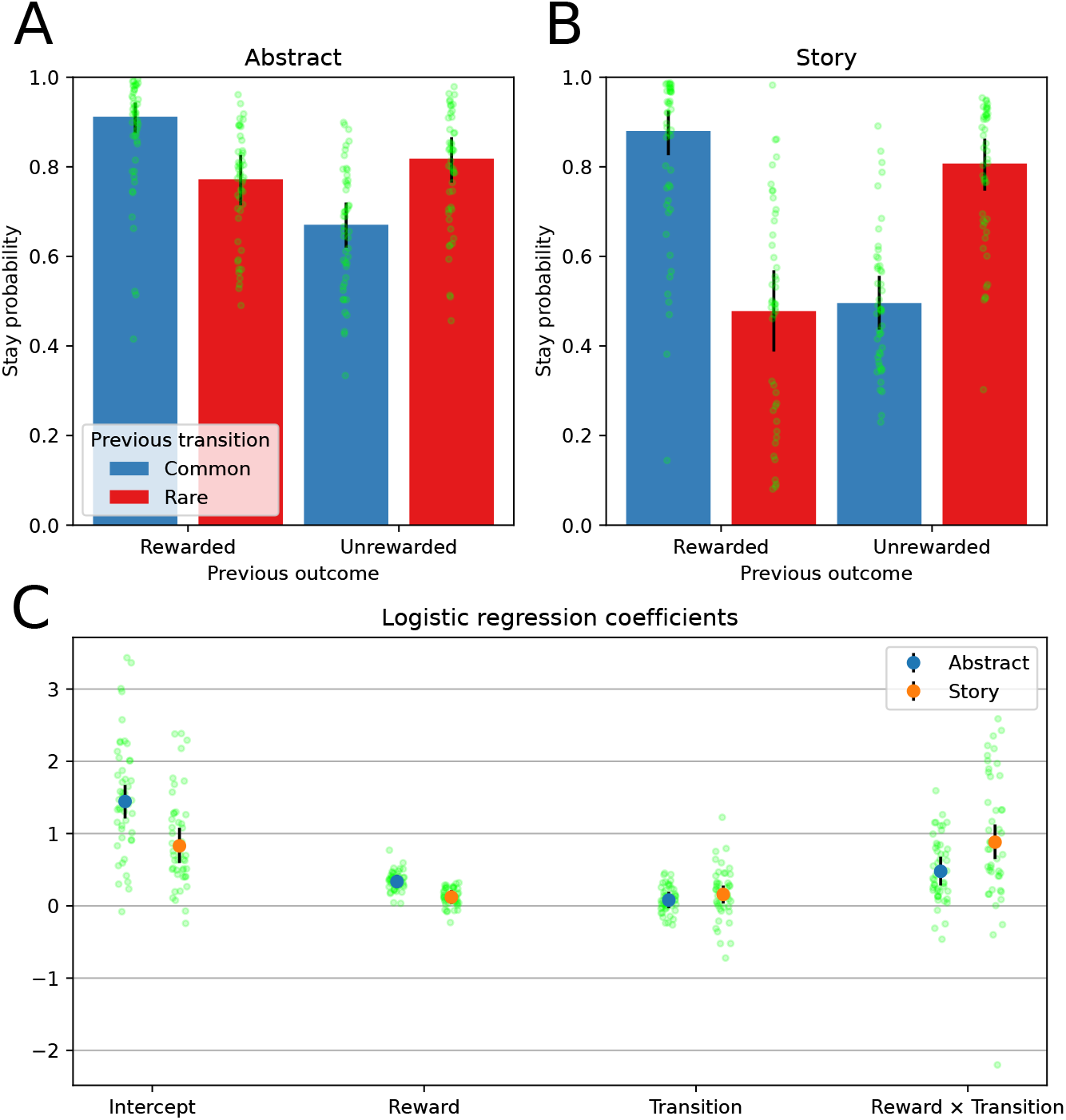
Results of a logistic regression analysis of the stay probability in consecutive trial pairs. A) Stay probabilities of the participants in the abstract condition (*N* = 48). B) Stay probabilities of the participants in the story condition (*N* = 46). C) Coefficients of the logistic regression model for participants in the abstract and story conditions. The story condition exhibiting a lower reward coefficient and a higher reward-by-transition coefficient compared to the abstract condition suggests that the former has adopted a more correct model-based strategy compared to the latter. These results replicate those of Feher da Silva and Hare [28]. Each bar height in subplots A and B and each blue or orange circle in subplot C represent the median value of the variable. Each green point represents the median value of the variable for each participant. Error bars represent the 95% high probability-density interval.

Next, we tested whether participants’ behaviour was better explained by this regression or a hybrid RL model. This hybrid RL model was proposed in the original paper [4] and is frequently used (e.g. [13, 3, 18, 26, 27, 24, 25]) to capture both learning and decision-making. In contrast to the logistic regression, it specifies the model-free and model-based strategies explicitly and does not allow for deviations aside from a weighted mixture of the two. Although there are many modifications of this model, they all explicitly define a limited set of algorithms an agent might employ. That is, they all make stronger assumptions than the logistic regression. We compared the two models to determine which one better explained participants’ first-stage choices. Model comparison showed that the logistic regression model is better at explaining first-stage choices than the hybrid model for both conditions. The difference in PSIS-LOO scores between the logistic regression and the hybrid models was −190.8 ±58.4 for the abstract condition and −1626.2 ±148.4 for the story condition. This replicates our previous findings [28].

We argue that if the hybrid model is a good approximation of behaviour or neural activity, it should explain the data at least as well as a simple regression model including only categorical regressors. However, it does not, which indicates that participants’ mental models of the task deviate substantially from its assumptions, and thus we should not draw strong conclusions from its fitted parameters. Nevertheless, we will use the hybrid RL model in some neuroimaging analyses when attempting to replicate past work. Therefore, we report the fitted parameters in the Supplemental Material.

#### Story instructions lead to better understanding and less effort

Next, we compared participants’ ratings for complexity, effort, and understanding across the two conditions (Figure 3). After performing the task, participants filled in a questionnaire in which they rated from 0 to 4 how well they understood the game (0: not at all, 4: perfectly), how complex they thought the game was (0: not at all, 4: extremely), and how effortful playing the game was (0: not at all, 4: extremely). We analyzed the ratings using ordered logistic models. Consistent with our hypothesis, participants in the abstract condition found the task more effortful than participants in the story condition (median: 3 in abstract, 2 in story, effect of story relative to abstract instructions on effort ratings: −0.42, 95% CI [−0.79, −0.05]; this effect is negative with 0.987 probability). The abstract group also reported lower understanding (median: 2 vs 3, effect of story relative to abstract instructions on understanding: 0.37, 95% CI [0.00, 0.76]; this effect is positive with 0.975 probability). On the other hand, although complexity and understanding ratings significantly correlated with one another (Spearman’s *ρ* = −0.419, 95% CI [−0.573 −0.265], *N* = 93, two-sided *P* < 0.001), complexity did not significantly differ between the two instruction groups (both medians: 2, effect of story relative to abstract instructions on complexity ratings: −0.12, 95% CI [−0.48, 0.25]; this effect is negative with 0.729 probability).

**Figure 3:**
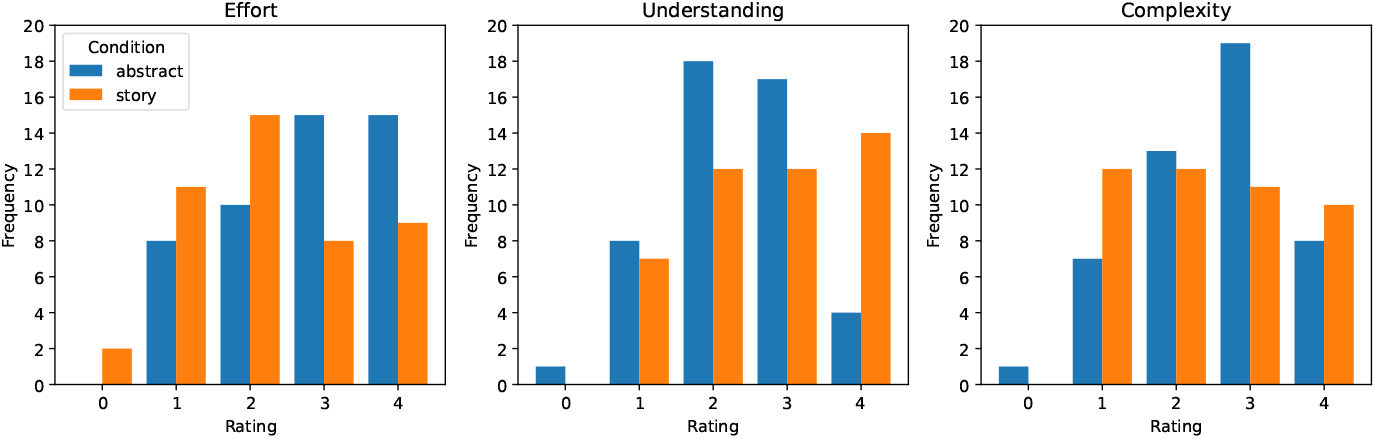
Distribution of ratings by condition for effort, understanding and complexity. After participants in the abstract (*N* = 48) and story (*N* = 46) conditions performed the two-stage task, they completed a questionnaire where they rated the task from 0 to 4 on effort, understanding, and complexity. Participants in the story condition gave significantly lower ratings for effort and higher ratings for understanding compared to participants in the abstract condition.

### Neuroimaging results

We analyzed the neuroimaging data to address three goals. First, we attempted to replicate previously reported correlations between BOLD activity and RPEs generated by the hybrid RL model, but ultimately discovered that the general linear model (GLM) used in previous studies contained two major flaws and should be revised. Next, we tested if, similar to choices, a GLM containing only categorical regressors for the task events explained BOLD activity in the region of interest—the nucleus accumbens—better than a GLM including RPEs derived from the hybrid RL model. We focused on the nucleus accumbens because this portion of the ventral striatum plays a central role in reward learning [34] and was used as the region of interest in the original paper [4]. Lastly, we tested if differences in brain activity between the abstract and story conditions are consistent with our hypothesis that the abstract group is expending more effort to understand the task and explore different mental models of its structure.

#### No evidence of correlations between second-stage model-free and model-based reward prediction errors and activity in the ventral striatum

To avoid potential differences due to instruction type, we used only participants who received the traditional abstract instructions (*N* = 48) when attempting to replicate previous fMRI results. Using the same GLM, we were able to replicate the previous results [4, 14, 26, 27, 31] that seemed to show correlations between model-based and model-free RPEs from the hybrid RL model and BOLD activity in the ventral striatum (Figure 4A). Surprisingly, we found that hybrid model parameters fit to the behavioural data from the original study [4] produced even higher correlations with our participants’ ventral striatal BOLD activity than model parameters fit to the behavioural data from our participants themselves (see Supplemental Material, Figure S2).

When running the replication analyses, we realized that the GLM used to explain the fMRI data in the original [4] and subsequent studies [14, 26, 27, 31] had problematic features, the most significant being that it combined each sequence of model-free or model-based RPEs at the second-stage with the sequence of RPEs at feedback into a single regressor (see Figure 1C for the timeline of a trial). Henceforth, we will refer to this model as the combined-RPE GLM. This model can only detect correlations between RPEs and the average of brain activity at the second stage and feedback, without distinction. However, it is possible that RPEs generated by the hybrid RL model correlate with brain activity at only one point, e.g., feedback, but not at the other. The combined-RPE GLM would not detect this situation. Critically, RPEs are easy to predict at feedback, but much more difficult at the second stage. At feedback, we can safely assume that when the participant received a reward, the RPE was positive and when they received no reward, the RPE was negative. It is unnecessary to combine model-based and model-free RL, or even to employ any specific RL theory, to make reasonable predictions about the RPE at feedback (we prove this in subsequent tests). However, it is much less clear what the RPE should be at the second stage. Here, different RL algorithms will make different predictions. The hybrid RL model posits that the RPE should be a combination of model-free and model-based signals. This is a strong prediction and to test it we must examine brain activity at the second stage alone.

We ran this test of the hybrid RL model by splitting the sequences of model-free and modelbased RPEs into separate regressors capturing signals at the second stage and at feedback individually. At the second stage, this separated-RPE GLM includes regressors for the model-free and model-based RPEs. At feedback, there is a single RPE regressor because the model-free and model-based RPEs are identical at that point. The results from this analysis show no statistically significant positive correlations between the model-free RPE and BOLD activity in any area (although there were negative correlations in some cortical areas). In particular, there were no significant correlations with activity in the nucleus accumbens, either at the whole brain level or with small volume correction (Figure 4B). The model-based RPE at the second stage correlated significantly with brain activity in several areas, including the medial prefrontal cortex and the left putamen, but not the nucleus accumbens. In short, neither model-free nor model-based RPEs from the hybrid RL model could explain activity in the nucleus accumbens at the crucial second stage, i.e., the only point in the task where model-free nor model-based RPEs are distinguishable.

**Figure 4:**
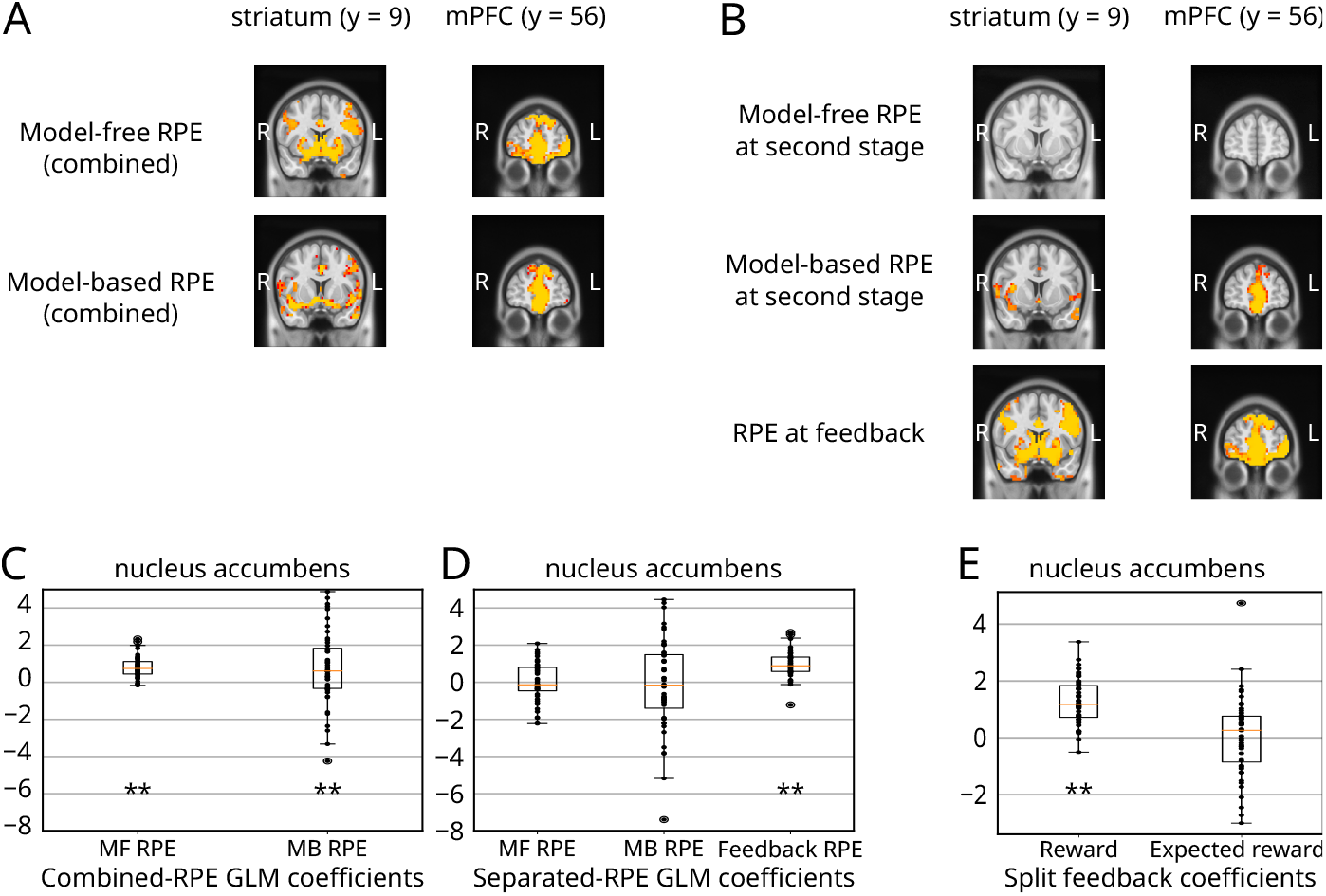
Correlation between BOLD activity and reward prediction errors (RPEs) in the striatum and the medial prefrontal cortex (mPFC) for the abstract condition (*N* = 48). Note that these results refer to the coefficients that describe BOLD activity as a linear function of the RPEs and not to the RPEs themselves. A) Combined-RPE GLM results, where regressors combine data from both the second stage and feedback. B) Separated-RPE model, which contains separate regressors for the second stage and feedback. C,D) Mean estimated coefficients from the combined-RPE (C) separated-RPE GLMs (D) within the nucleus accumbens. E) Mean estimated coefficients from the two components of the feedback RPE—reward and expected reward—within the nucleus accumbens. They were obtained by splitting the feedback RPE from the combined-RPE GLM into reward and expected reward, the feedback RPE being the difference between these values. In subplots A–B, voxelwise analyses were performed at the whole brain level with a family-wise-error-corrected threshold of *P* = 0.05. In subplots C–E, each black dot represents the mean coefficient from a single participant. The box and whisker plots show the distribution across the entire sample. The box extends from the first quartile to the third quartile of the distribution, with a line at the median. The whiskers extend from the box by 1.5 times the inter-quartile range. The symbol ** indicates statistically significant results as determined by one-sample one-sided Wilcoxon signed-rank tests (*N* = 48). The associated p-values from left to right, corrected for multiple comparisons, are < 0.001, 9.90 × 10^-3^, 1.00, 1.00, < 0.001, < 0.001, 1.00, with the corresponding *V* values being 1162, 862, 596, 574, 1141, 1167, 614.

Conversely, the results at feedback were very similar to those obtained for the model-free RPE regressor in the combined-RPE GLM. This suggests that the combined-RPE GLM results for the model-free RPE were due to its correlation with brain activity at feedback, not at the second stage. Indeed, plotting the average coefficients in the nucleus accumbens from the combined and separated-RPE GLMs at each stage shows exactly this pattern (Extended Data Figure 1). These findings were surprising, so we investigated further. We conducted mathematical analyses that revealed that the apparent correlation between hybrid model RPEs and brain activity in the combined-RPE GLM is overestimated and most likely spurious given the results from the separated-RPE GLM (see Supplemental Material).

The shortcomings of the combined-RPE GLM and our results from the separated-RPE GLM imply that neither our current data nor any previous publications provide reliable evidence of model-free RPEs in the nucleus accumbens during the two-stage task. It is possible that analyzing other data sets with the separated-GLM will show model-free RPEs at the second stage. However, given current and previous behavioural results [22, 28] and the aggregate fMRI data to date, it seems likely that the simple model-free learning algorithm assumed by the hybrid RL model plays little role in the two-stage task.

#### Basic categorical regressors explain activity in the nucleus accumbens better than hybrid-model-derived RPEs

Next, we tested if, consistent with the behavioural results, a basic categorical GLM explained activity in ventral striatum better than the two GLMs including RPEs from the hybrid RL model. The categorical GLM contained the first-stage parametric modulators derived from the hybrid model to isolate the comparisons to the second stage and feedback (Figure 1C). Two separate model comparison metrics favored the categorical GLM over the other GLMs (Table 1). These results indicate that categorical task events explain activity in the nucleus accumbens better than RPEs from the hybrid RL model. Thus, neither choice behaviour nor nucleus accumbens BOLD signals are best-explained by a hybrid RL model that assumes participants only use a combination of simple model-free and correct model-based strategies.

**Table 1:**
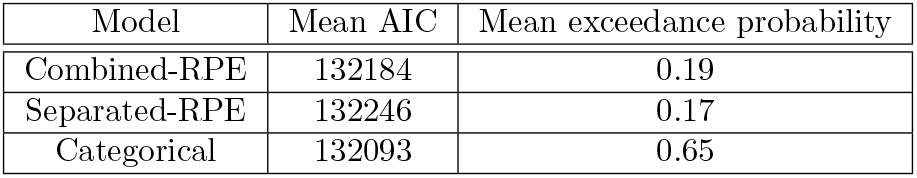
Model comparison between fMRI models in the nucleus accumbens (*N* = 48). Three fMRI models (combined-RPE, separated-RPE, and categorical) were used to correlate BOLD activity in the nucleus accumbens to different predictors. Both the AIC and the mean exceedance probability were calculated for each model. The best model according to both analyses (with the smallest AIC score and the largest mean exceedance probability) was the categorical model, which had two binary predictors: the transition (common or rare) and the outcome (reward or no reward) for each trial.

| Model | Mean AIC | Mean exceedance probability |
| --- | --- | --- |
| Combined-RPE | 132184 | 0.19 |
| Separated-RPE | 132246 | 0.17 |
| Categorical | 132093 | 0.65 |

The categorical GLM revealed that several brain regions respond differentially to common versus rare transitions or rewarded versus unrewarded outcomes. This model included separate onset regressors for the second-stage and feedback as well as categorical modulators at the second stage, indicating transition (common = +1, rare = −1), and feedback, indicating outcome (reward = +1, no reward = −1). The transition regressor encodes a state prediction error (SPE; [35]) but is also a good proxy for model-agnostic RPEs at the second stage. Although SPEs and RPEs are conceptually distinct, they are correlated in the two-stage task, assuming that the common transition takes the participant to their most desired state. However, similar to the separated-RPE GLM results, the transition regressor was not significantly correlated with activity in any voxels within the nucleus accumbens, although many other brain areas were sensitive to the transition type (see Supplemental Table S3).

Conversely, the outcome regressor was positively correlated with activity in all voxels within the nucleus accumbens. These results are similar to those of the feedback RPE from the separated-RPE GLM. The feedback RPE is defined as the reward minus the expected reward, therefore we tested whether the expected reward, which is the only term derived from the hybrid RL model, was significant after accounting for the reward. Specifically, we split the feedback RPE from the separated-RPE model into reward and expected reward regressors and fit this model to the data. Figure 4E, shows that only reward is significant.

#### Abstract instructions lead to more brain activity at the second stage and feedback

We pursued our third aim by analyzing the differences in BOLD activity between the abstract and story conditions. We focus on the feedback stage because reward learning theory holds that updating should occur then and previous findings [19] indicate that decisions may be made at this point as well (see Supplemental Material for the first- and second-stage results). At feedback, participants in the abstract relative to the story condition exhibited higher activity in regions including the right inferior prefrontal cortex, the right frontopolar cortex, the ventral anterior insula, and the right caudate (Figure 5). The bilateral inferior frontal cortex and right frontopolar cortex were previously found to encode the reliability of model-based and model-free strategies [36]. The right frontopolar cortex was also found to encode the reliability of different strategies in a pattern learning task [37], and the right ventrolateral prefrontal cortex was found to be involved in the arbitration between choice imitation and goal emulation [38]. Moreover, the right frontopolar cortex has been linked to exploratory decisions [39, 40]. Although we should draw inferences about behaviour or mental states from brain activity with caution, these results are consistent with our hypothesis that participants in the abstract condition have less certainty about their task model and update their conceptualization of it after feedback. These data also accord with reports of less understanding and more effort in the abstract compared to story condition.

**Figure 5:**
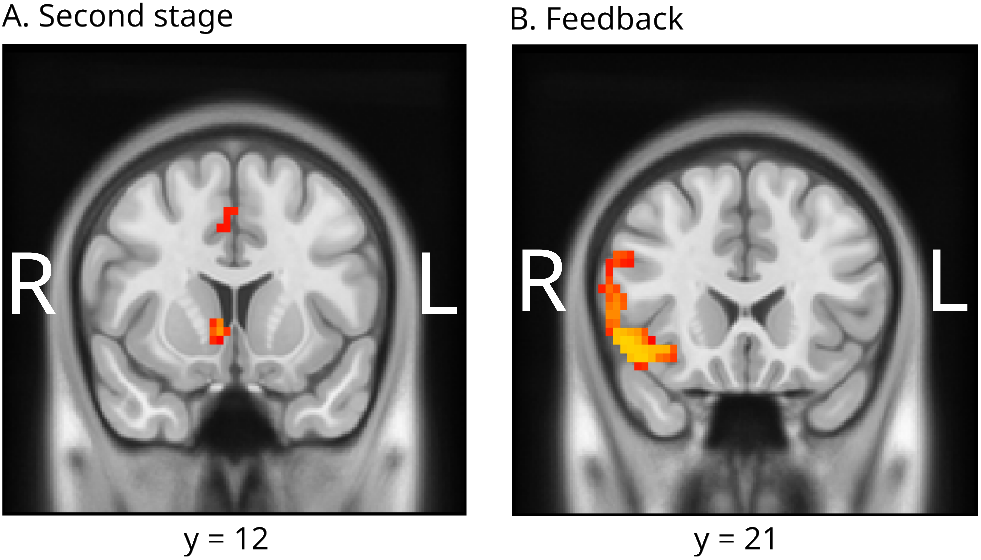
Brain activity comparison between the abstract (*N* = 48) and story (*N* = 46) conditions at A) the second stage and B) feedback. No differences above the threshold were found between the two conditions at the first stage. All analyses were performed at the whole brain level with a family-wise-error-corrected threshold of *P* = 0.05. See more results in the Supplemental Material.

**Figure 6:**
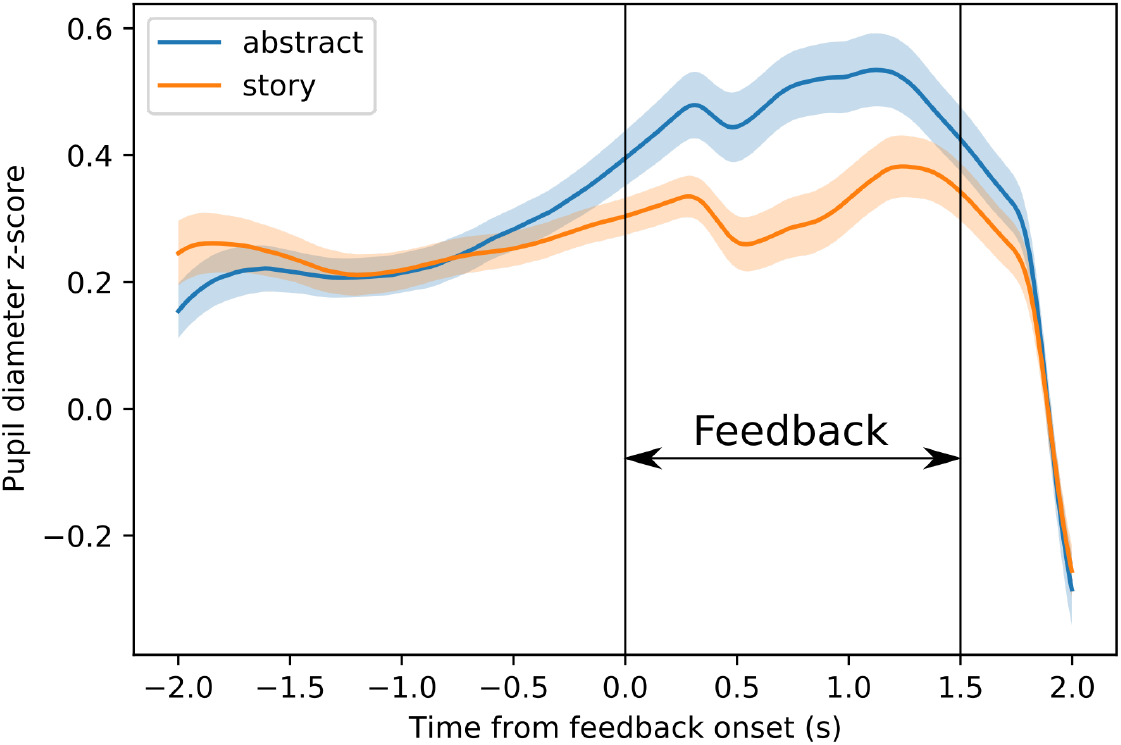
Pupil diameter z-scores at feedback for the abstract (*N* = 36) and story (*N* = 37) conditions. Our analysis found that participants in the abstract condition had a higher pupil diameter at feedback compared to participants in the story condition.

#### Participants in the abstract condition have larger pupil diameter at feedback

Larger pupil diameter has been linked to mental effort [41, 42], exploration [43], arousal [44], uncertainty [45, 46], surprise [47], and cognitive control [48]. Although we cannot infer the exact causes of pupil diameter changes in the two stage task, these previous reports suggest that pupil diameter should be larger after feedback if there is more effort or less understanding. Thus, we predicted that participants in the abstract condition would exhibit a larger pupil diameter after feedback.

We examined pupil diameter at feedback with a mixed-effects linear model including as predictors the condition, reward, and transition (all encoded as −1 and +1) as well as all their interactions. None of the interactions were statistically significant (see Supplemental Material). There were main effects of condition (story < abstract, −0.09 95% CI [−0.15, −0.02]), reward (reward > no reward, 0.05 95% CI [0.03, 0.08]), and transition (rare > common, 0.07 95% CI [0.05, 0.10]).

## Discussion

We hypothesized that participants who seemingly employ a hybrid model-free/model-based strategy in the two-stage task are in fact exploring different incorrect model-based strategies, which makes performance more effortful than using a single model-based strategy. Our behavioural, fMRI, and pupil diameter data all support this hypothesis. Replicating our previous results [28], participants in the story condition displayed nearly correct model-based behaviour in the two-stage task, whereas participants in the abstract condition seemingly employed a hybrid strategy. However, participants in the abstract condition rated the task as more effortful compared to the story condition. During feedback, participants in the abstract condition had larger pupil diameters, which have been linked to exploration [43] and mental effort [41, 42], and exhibited greater activity in prefrontal regions previously linked to strategy switching and arbitration [37, 36]. We must make inferences about mental states from physiological data with caution, but overall it seems that the participants in the abstract condition, who appeared to rely more on model-free learning, did not reap the savings in effort that model-free behaviour provides. Rather, the evidence is more consistent with our hypothesis that participants in the abstract condition frequently arbitrated between incorrect mental models of the task structure.

Our results add to the set of findings that appear to contradict the idea that participants will only use a model-based strategy when it pays off more than model-free learning. Recall that in the two-stage task, the model-based strategy does not pay off more and is more computationally costly than the simple model-free strategy [3, 49, 50]. Therefore, an agent combining model-free and model-based strategies should become more model-free as it learns that performing effortful model-based calculations does not lead to greater payoffs. Yet, contrary to this prediction, participants maintain their behaviour [25] or even become more model-based as they practice the two-stage task [13]. One potential explanation for these findings is that participants may employ multiple strategies beyond the correct model-based and simple model-free ones, e.g., sophisticated model-free algorithms or goal-direct hierarchies of action sequences [6, 2, 51, 28, 52, 53]. The proposal that participants deviate from the assumed strategies is supported by our findings that the hybrid algorithm fails to explain both choice data and BOLD signals in the ventral striatum when compared to simple categorical models.

It is also possible that participants save effort by not employing the full model-based strategy and using an approximation instead. For example, as each trial starts, participant may think about the second-stage choice that they believe has the highest reward probability and then just select the first-stage action that is most likely to transition to that state (i.e., they prune the branches of the decision tree that are less likely to happen—the rare transitions). This is not the full model-based strategy: to calculate the model-based value of each first-stage option, one must take into account both common and rare transitions. However, each first-stage option is more likely to lead to a different second-stage state, and transition probabilities are the same for both first-stage options. Thus, the heuristic above is behaviourally indistinguishable from the full model-based strategy. Participants using it will appear model-based to common analyses but in reality will exert less effort than the full model-based strategy requires. Unfortunately, it is impossible to determine if participants use the full or this simplified model-based strategy by analyzing their choices. Beyond these specific possibilities, there are innumerable reinforcement-learning strategies and heuristics that all come with different effort-reward trade-offs.

Critically, none of these alternative strategies increase payoffs or reduce effort in the two-stage task compared to a simple model-free strategy [2, 3]. Why would participants converge to them as opposed to a simple model-free strategy? To answer this question, we must consider that simple model-free learning may not be automatic or distinct from model-based or heuristic strategies at the neural level. Thus, participants may never use a model-free strategy in the two-stage task, either because it is not congruent with their mental model of the task or for other reasons.

Consistent with this idea, we did not find any brain regions where BOLD activity positively correlated with the model-free RPEs at the second stage when employing a direct test for these signals (the separate-RPE GLM). Previous reports that both model-free and model-based RPEs were significantly correlated with BOLD activity in several brain areas in the two stage task [4, 14, 26, 27], as well as three other studies reporting similar results from a related multi-stage decision task [35, 36, 54], were based on an analysis that did not make the critical distinction between the second stage and feedback. In particular, the neuroimaging analyses in the original two-stage task paper were aimed at testing if adding a model-based RPE regressor to a model that already included a model-free one would improve the model’s fit to neural activity in the ventral striatum and the medial prefrontal cortex [4]. Following the general opinion in the field, the authors took it as given that model-free RPEs would correlate with neural activity in those areas. However, the distinction between RPEs at the second stage and feedback is essential in order to test the hybrid hypothesis because the two strategies produce different RPEs only at the second stage.

Critically, when we tested the predictions of the hybrid hypothesis using an fMRI model that distinguished between second stage and feedback, we did not find evidence of either model-free or model-based RPEs in the nucleus accumbens at the second stage. Ultimately, the model that best predicted activity in the nucleus accumbens was a simple categorical model with a regressor for common and rare transitions at the second stage and a regressor for reward and no reward at feedback. This result parallels the behavioural findings that a simple logistic regression model fit the first-stage choice data better than the hybrid reinforcement learning model.

The categorical models of behaviour and BOLD activity are simplistic approximations of neural processes. We do not claim they are better than the hybrid model at providing mechanistic insights. Rather, we argue that if we aim to draw robust inferences from a model of the potential mechanisms that underlie behaviour in a multi-stage decision task, then such a model should explain the data as well as or better than a simplistic categorical model. Moreover, we acknowledge that, in general, striatal activity correlates with model-based RPEs when such strategies are used; this has been repeated using a variety of methods across species [55, 56, 57]. Rather, we challenge the hybrid hypothesis that both model-free and model-based RPEs are computed in parallel and have a combined influence on the ventral striatum. To our knowledge, all studies that have shown evidence for this employed multi-stage tasks performed by human participants in an fMRI scanner and did not make the crucial distinction between different task stages in their analyses.

Although many brain regions were sensitive to common versus rare transitions to the second stage, we did not find significant differences in the nucleus accumbens. Moreover, nucleus accum-bens activity did not significantly correlate with either simple model-free or correct model-based RPEs. In contrast, there was much stronger activity in the nucleus accumbens for reward receipt compared to omission. These results could be due to the reward probabilities of the second-stage options drifting over time, which left participants unsure about how rewarding each second-stage state is. Consequently, arriving at the second stage may generate only small RPEs. Moreover, participants may employ diverse strategies, which lead to highly variable RPEs that don’t aggregate to yield a consistent pattern at the group level. In addition, the original task design, which we used to facilitate comparisons with previous studies, did not include a temporal jitter in the interval between the second stage and feedback except for small variations in response times (approximately 300 ms). This may limit the ability to distinguish between second-stage and feedback signals in the present and previous studies.

Overall, our findings contradict the idea of an automatic model-free system that is neurobiologically distinct from a model-based system. In addition, our findings raise strong doubts about the validity of previous reports of combined model-free and model-based RPEs in the ventral striatum. Regardless of the algorithm under investigation, our findings indicate that future work on reward learning and mental effort should explicitly include the cost of switching between candidate strategies in addition to the costs of executing the strategies to better understand decision-makers’ sensitivity to cost-benefit trade-offs. Often, a complex but stable strategy may be less effortful than repeatedly searching through potential strategies of varying complexity.

## Methods

### Sample size

One hundred human participants were invited to participate in this experiment. The sample size was determined by a power analysis (*R* = 0.4, *β* = 0.8, two-tailed *P*-value = 0.05, *N* required = 48) based on our previous behavioural findings [28]. These 100 participants were randomly assigned to the story (N = 50) and abstract conditions (N = 50). Data collection and analysis were not performed blind to the conditions of the experiments. This study was preregistered on 22 Dec 2018 [58].

Participants between 18–30 years old were recruited from the University of Zurich’s Registration Center for Study Participants. The experiment was conducted in accordance with the Zurich Cantonal Ethics Commission’s norms for conducting research with human participants, and all participants provided written informed consent on the study day. Participants had normal or corrected-to-normal vision. They were instructed not to take any medication or alcohol for 24 hours prior to the study and to come into the lab without make-up or remove it on site to prevent metallic components of the make-up from being moved or heated by the MRI machine. For the list of inclusion and exclusion criteria used in this study, see the Supplemental Material. The exclusion criteria included requirements for a food decision-making task the participants performed after the experiment described in this paper was over. Moreover, because of the food task, participants were instructed to eat a light meal 3 hours before the session (no more than 400 calories) and then for 2.5 hours consume nothing but water.

Out of the 100 participants who were invited to participate in the experiment, we excluded 4 participants assigned to the story condition (final *N* = 46) and 2 participants assigned to the abstract condition (final *N* = 48). The reasons for exclusion were as follows. One participant was excluded because the task presentation program crashed during the experiment and some behavioural results were lost, two participants received the wrong instructions (i.e., those meant for participants in the other condition), one participant fell asleep in the fMRI scanner during the task, one participant could not complete the required pregnancy test before entering the fMRI scanner, and one participant’s fMRI images were grossly distorted as determined by visual inspection. Questionnaires were completed by 93 of the final 94 participants (*N* = 48 in the abstract condition, *N* = 45 in the story condition).

### Task performance

Outside the fMRI scanner, participants received instructions for the practice task, which were different depending on the condition they were randomly assigned to. Participants in the story condition also completed the same quiz for understanding that we used in previous work [28]. All participants then performed 50 practice trials and read the instructions for the fMRI task. Inside the fMRI scanner, participants performed 150 trials of the two-stage task in either the abstract or the story condition. The task was performed over three fMRI runs, each containing 50 trials and lasting 842.5 seconds (around 14 minutes). Brain images were acquired using a 3T Philips Achieva scanner and preprocessed with fMRIprep 1.2.5 ([59, 60]; RRID:SCR_016216), as detailed below.

For the two-stage task, we recorded each participant’s decisions and reaction times. We also recorded neural activity, eye movements, and pupil diameter by using an eye tracker while the participant performed the task inside the fMRI scanner. Participants then completed a questionnaire where they were asked to describe the strategy they used to make choices, how well they understood the task, and how complex and effortful they thought the task was.

Functional imaging was performed using a 3T Philips Achieva scanner with an eight-channel sensitivity-encoding head coil (Philips Medical Systems) to acquire gradient echo T2*-weighted echo-planar images with BOLD contrast. Forty-two slices were acquired in ascending order with 3mm thickness, 3 × 3mm^2^ voxel resolution, 0.6mm gap between slices, repetition time of 2.5s, echo time of 30ms, and a flip angle of 90°. Participants were placed in a cushioned head restraint within the head coil to limit head movement. T1-weighted structural images, electrocardiograms, and respiratory measurements were also acquired.

### One-trial-back logistic regression analysis

This analysis employed a hierarchical logistic regression model whose parameters were estimated through Bayesian computational methods. The predicted variable was *p*_stay_, the stay probability, and the predictors were *x_r_*, which indicated whether a reward was received or not in the previous trial (+1 if the previous trial was rewarded, −1 otherwise), *x_t_*, which indicated whether the transition in the previous trial was common or rare (+1 if it was common, −1 if it was rare), the interaction of the two. Thus, for each participant in both conditions, an intercept *β*_0_ and three coefficients were determined, as shown in the following equation:

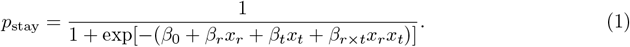

For each trial pair, the variable *y* was equal to 1 if the participant chose in the next trial the same first-stage action as in the previous trial (a “stay” choice) or equal to 0 if the participant chose a different first-stage action (a “switch” choice). The distribution of y was Bernoulli(*p*_stay_).

The model was fitted to all participants on both conditions with a Bayesian hierarchical model. The distribution of the 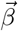 vectors was multivariate Student-t distribution with *v* = 4 degrees of freedom, for more robustness, rather than multivariate normal. The mean of this distribution 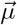 was given by a linear model with the participant’s condition *d* (abstract encoded as 0, story encoded as 1) as a predictor: 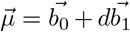, where 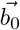 and 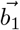 are coefficient vectors given the prior distributions 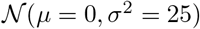 and Cauchy(0, 1) respectively. The covariance matrix **Σ** of the distribution was decomposed into a diagonal matrix *τ*, whose diagonal components were given Cauchy(0,1) prior, and a correlation matrix **Ω**, which was given an LKJ prior [61] with shape *v* = 2 [62], so that **Σ** = *τ***Ω**.

To obtain parameter estimates from the model’s posterior distribution, we coded the model into the Stan modeling language [63, 62] and used the PyStan Python package [64] to obtain 64 000 samples of the joint posterior distribution from four chains of length 32000 (warmup 16 000). Convergence of the chains was indicated by 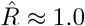 for all parameters.

### The hybrid reinforcement learning algorithm

This algorithm combines the model-free SARSA(λ) algorithm with model-based learning to perform the two-stage task [4]. At the start of the first trial *t* =1, the model-free values 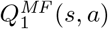 of each action *a* that can be performed at each state s are equal to zero. At the end of each trial t, the model-free values of the chosen actions are updated based on RPEs. For an action *a*_2_ performed at the second-stage state *s*_2_, the RPE is defined as 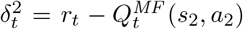, i.e. as the difference between the obtained reward *r_t_* and *a*_2_’s current value. Then *a*_2_’s model-free value is updated as follows:

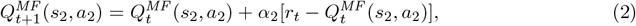

where *α*_2_ is the second-stage learning rate (0 ≤ *α*_2_ ≤ 1). For an action *a*_1_ performed at the first-stage state *s*_1_, the reward prediction error is defined as 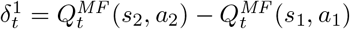, but _a1_’s model-free value is updated based on both the first and second stages, as follows:

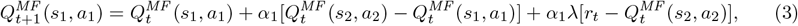

where *α*_1_ is the first-stage learning rate (0 ≤ *α*_1_ ≤ 1), and *λ* is the eligibility parameter (0 ≤ *λ* ≤ 1), which modulates the effect of the second-stage RPE on the values of first-stage actions.

The model-based value 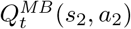 of each action *a*_2_ performed at second-stage state *s*_2_ is the same as its model-free value:

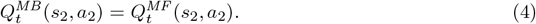

The model-based value of each first-stage action *a*_1_ is calculated (rather than cached, as the model-free value) from the values of second-stage actions as follows:

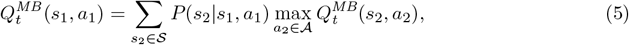

where *P*(*s*_2_|*s*_1_, *a*_1_) is the probability of transitioning to second-stage state *s*_2_ by performing action *a*_1_ at first-stage state *s*_1_, 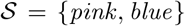 is the set of second-stage states, and 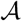 is the set of actions available at that state.

First-stage choices are made based on both the model-free and the model-based values of each first-stage action, weighted by a model-based weight *w* (0 ≤ *w* ≤ 1), according to a soft-max distribution:

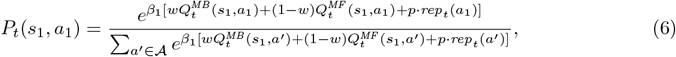

where *β*_1_ is the first-stage’s inverse temperature parameter, which determines the exploration rate at this stage, *p* is a perseveration parameter that add a tendency to repeat the previous trial’s first-stage action in the next trial, and *rep_t_*(*a*′) is equal to one if the agent performed the first-stage action *a*′ in the previous trial, and to zero otherwise. At the second stage, the model-free and model-based values of second-stage actions are the same and there is no assumed tendency to repeat the previous action. Second-stage choice probabilities are given by:

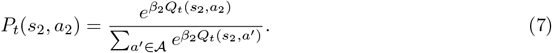

The hybrid model was fitted to all participants on both conditions with a Bayesian hierarchical model, which allowed us to pool data from all participants to improve individual parameter estimates and easily calculate the condition effect on each parameter.

The parameters of the hybrid model for the *i*th participant were 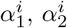, *λ^i^*, 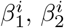, *w^i^*, and *p^i^*. Vectors

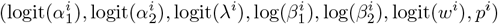

were generated for each participant, and each component of the vectors was given a Student-t distribution with *ν* = 4 degrees of freedom. These transformations of the parameters were used because many of the original values were constrained to an interval and the transformed ones were unbounded, which is required by the Student-t distribution. The mean *μ* of this distribution was given by *μ* = *b*_0_ + *b*_1_*d*, where *d* was the participant’s condition d (abstract encoded as 0, story encoded as 1). The standard deviation of this distribution as well as the coefficients *b*_0_ and *b*_1_ were given Cauchy(0, *σ* = 5) priors.

This hierarchical model was coded in the Stan modelling language [62, 63] and fitted to our data set through the PyStan interface [64] to obtain two chains of 32000 iterations (warmup: 16 000). Convergence was indicated by 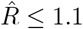 for all parameters.

We fitted the same model to our data from the abstract condition by maximum likelihood, using 1,000 applications of Stan’s optimization algorithm, and allowing a different *w* for each participant, but keeping all the other parameters the same for all participants. Additionally, we fitted the model separately for each participant by maximum likelihood, using the same method as above, and calculated the mean parameters for each condition.

### Behavioural model comparison

The PSIS-LOO IC is an estimate of the out-of-sample prediction error, and smaller values indicate better performance [65]. To obtain PSIS-LOO (an approximation of leave-one-out cross-validation) scores, the log-likelihood of every trial was calculated for each iteration and used as input to the `loo` and `compare` functions of the loo R package [65].

Note that the hybrid model was fitted to both first-stage and second-stage choices, but because the logistic regression model can only be fitted to first-stage choices, model comparison scores were computed only for first-stage choices. At first, it may seem more fair to also fit the hybrid model only to first-stage choices. However, the two stages are not independent. The hybrid model uses the second-stage choices to calculate the second-stage action values, and from them it calculates the first-stage action values and predicts the first-stage choices. Hence, it only makes sense to fit this model to both stages.

### Questionnaire ratings analysis

Ratings for effort, understanding, and complexity were modelled with ordered logistic distributions separately for each rating category [66]. Each distribution had two parameters: 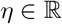, which was given a weak 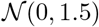 prior, and 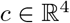 such that *c_k_* < *c*_*k*+_ for *k* ∈ 1, 2, 3, each element of which was given a weak 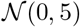 prior (the ordering constraint on *c* truncates the joint prior of its elements to points that satisfy this constraint). For each participant, the analysis took into account their rating *r* ∈ 0, 1,&, 4 and condition *d*, encoded as −1 if abstract and 1 if story. The likelihood of each participant’s rating *P*(*r, d*) was given by

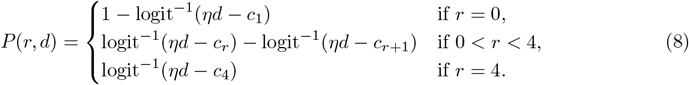

Thus, η was the effect of condition on the ratings. If *η* > 0, then ratings were larger for the story condition and if *η* < 0, then ratings were larger for the abstract condition. This model was coded on Stan and, for each rating category, four chains were run for 12 000 iterations (2 000 warm-up). Convergence of the chains was indicated by 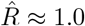 for all parameters. The minimum number of effective samples was 15 000.

### Preprocessing of fMRI images

Results included in this manuscript come from preprocessing performed using fMRIPrep 1.2.5 ([59, 60]; RRID:SCR_016216), which is based on Nipype 1.1.6 ([67, 68]; RRID:SCR_002502).

#### Anatomical data preprocessing

The T1-weighted (T1w) image was corrected for intensity non-uniformity (INU) using `N4BiasFieldCorrection` [69, ANTs 2.2.0], and used as T1w-reference throughout the workflow. The T1w-reference was then skull-stripped using `antsBrainExtraction.sh` (ANTs 2.2.0), using OASIS as target template. Spatial normalization to the ICBM 152 Nonlinear Asymmetrical template version 2009c [70, RRID:SCR_008796] was performed through nonlinear registration with `antsRegistration` [ANTs 2.2.0, RRID:SCR_004757, 71], using brain-extracted versions of both T1w volume and template. Brain tissue segmentation of cerebrospinal fluid (CSF), white-matter (WM) and gray-matter (GM) was performed on the brain-extracted T1w using `fast` [FSL 5.0.9, RRID:SCR_002823, 72].

#### Functional data preprocessing

For each of the 3 BOLD runs found per subject (across all tasks and sessions), the following preprocessing was performed. First, a reference volume and its skull-stripped version were generated using a custom methodology of fMRIPrep. A deformation field to correct for susceptibility distortions was estimated based on fMRIPrep’s fieldmap-less approach. The deformation field is that resulting from co-registering the BOLD reference to the same-subject T1w-reference with its intensity inverted [73, 74]. Registration is performed with `antsRegistration` (ANTs 2.2.0), and the process regularized by constraining deformation to be nonzero only along the phase-encoding direction, and modulated with an average fieldmap template [75]. Based on the estimated susceptibility distortion, an unwarped BOLD reference was calculated for a more accurate co-registration with the anatomical reference. The BOLD reference was then co-registered to the T1w reference using `flirt` [FSL 5.0.9, 76] with the boundary-based registration [77] cost-function. Coregistration was configured with nine degrees of freedom to account for distortions remaining in the BOLD reference. Head-motion parameters with respect to the BOLD reference (transformation matrices, and six corresponding rotation and translation parameters) are estimated before any spatiotemporal filtering using `mcflirt` [FSL 5.0.9, 78]. BOLD runs were slicetime corrected using `3dTshift` from AFNI 20160207 [79, RRID:SCR_005927]. The BOLD time-series (including slice-timing correction when applied) were resampled onto their original, native space by applying a single, composite transform to correct for head-motion and susceptibility distortions. These resampled BOLD time-series will be referred to as preprocessed BOLD in original space, or just preprocessed BOLD. First, a reference volume and its skull-stripped version were generated using a custom methodology of fMRIPrep. Several confounding time-series were calculated based on the preprocessed BOLD: framewise displacement (FD), DVARS and three region-wise global signals. FD and DVARS are calculated for each functional run, both using their implementations in Nipype [following the definitions by 80]. The three global signals are extracted within the CSF, the WM, and the whole-brain masks. Additionally, a set of physiological regressors were extracted to allow for component-based noise correction [CompCor, 81]. Principal components are estimated after high-pass filtering the preprocessed BOLD time-series (using a discrete cosine filter with 128s cut-off) for the two CompCor variants: temporal (tCompCor) and anatomical (aCompCor). Six tCompCor components are then calculated from the top 5% variable voxels within a mask covering the subcortical regions. This subcortical mask is obtained by heavily eroding the brain mask, which ensures it does not include cortical GM regions. For aCompCor, six components are calculated within the intersection of the aforementioned mask and the union of CSF and WM masks calculated in T1w space, after their projection to the native space of each functional run (using the inverse BOLD-to-T1w transformation). The head-motion estimates calculated in the correction step were also placed within the corresponding confounds file. All resamplings can be performed with a single interpolation step by composing all the pertinent transformations (i.e. head-motion transform matrices, susceptibility distortion correction when available, and co-registrations to anatomical and template spaces). Gridded (volumetric) resamplings were performed using `antsApplyTransforms` (ANTs), configured with Lanczos interpolation to minimize the smoothing effects of other kernels [82]. Non-gridded (surface) resamplings were performed using `mri_vol2surf` (FreeSurfer).

Many internal operations of fMRIPrep use Nilearn 0.5.0 [83, RRID:SCR_001362], mostly within the functional processing workflow. For more details of the pipeline, see the section corresponding to workflows in fMRIPrep’s documentation.

After images were preprocessed with fMRIPrep, preprocessed images were smoothed using a Gaussian kernel with full-width at half-maximum (FWHM) of 6 mm.

### Exclusion of runs and volumes

Movements over 3 mm across consecutive volumes (i.e. framewise displacement larger than 3 mm) led to the exclusion of the affected volume. To ensure movement artefacts were fully excluded, the volume before and three volumes after the affected scan were excluded from the main analysis. We also excluded volumes marked as “non-steady-state” by fmriprep as well as volumes marked as “motion outliers” (having framewise displacement greater than 0.5 mm or standardised DVARS greater than 1.5 mm). Functional runs where over 30% of TRs were affected were entirely excluded. In total, six runs were excluded, two from the same participant and four from another four participants.

### First-level analyses

To compare brain activity between the story and abstract conditions, we used a first-level model with three onset regressors (O1, O2, O3) for the first stage, second stage, and feedback, respectively. The duration of O1 and O2 was set equal to the reaction time, and the duration of O3 was set equal to feedback duration (1.5 s). Regressors O1 and O2 had the reaction time as a parametric modulator. This model did not contain any RPE regressors. Regressors for this model and the models described in the next paragraphs were mean-centered and orthogonalized following SPM12’s default procedure.

To replicate the results from the original study [4], we estimated their original, combined-RPE GLM, which included the model-free RPE and the difference between the model-based and model-free RPEs as parametric regressors of impulse events (of zero duration) at the second stage and feedback. The combined-RPE GLM also included nuisance onsets at the first stage and at feedback, and the first stage regressor was modulated by two parametric regressors: the value of the chosen first-stage action and its partial derivative with respect to w. The model-based RPE was defined as

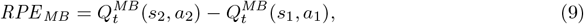

with the value of 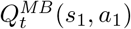 given by Equation 5, and the model-free RPE was defined as

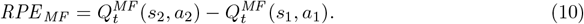

Note that 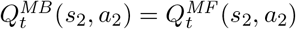. The RPE at feedback was defined as

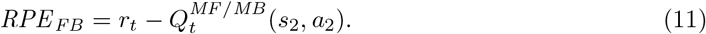

The separated-RPE GLM we introduce in this work was identical to the original combined-RPE GLM at the first stage, but split the second stage and feedback events into separate regressors. The separated-RPE GLM split also the model-free regressor into two regressors, separating second-stage events from feedback events. The difference in MB minus MF RPEs was only included for the second-stage events, because this difference was always zero at the feedback events. We used the hybrid parameters from the original study [4] to calculate the RPE for this model, because they generated the strongest correlations with brain activity in the combined-RPE GLM.

The categorical GLM included categorical modulators for transition (1 for common and −1 for rare) at the second stage and for reward (1 for reward and −1 for no reward) at feedback. It did not contain any RPE regressors.

The first-level analyses we conducted were restricted to a grey matter mask made from the conjunction of all grey matter masks created by fmriprep for all participants. First-level analyses were performed using SPM12. To correct physiological artefacts created by heart rate and breathing, the default pipeline of the PhysIO toolbox was employed to obtain 18 regressors for each run. Additionally, to correct artefacts caused by head motion, and following a common recommendation for fmriprep [84], we added to the model six regressors for rotation and translation, a regressor for the framewise displacement, and the six noise regressors calculated using anatomical CompCor.

### Second-level analyses

Second-level analyses were performed using FLS’s randomise tool. The algorithm was run to generate 5 000 permutations of the data with Threshold-Free Cluster Enhancement (TFCE) and 6 mm variance smoothing for t-stats. The number of runs performed by each participant was added as a nuisance variable to all analyses. In second-level analyses of contrasts concerning model-based RPEs, the model-based weight was also added as a variable, following the original study [4]. The model-based weight was obtained by maximum likelihood estimation given the other hybrid parameter values used in the analysis, i.e. our mixed-effects model estimates or the mixed-effects estimates from the original paper [4]. The number of runs per participant was added as a nuisance variable to account for participants with excluded runs. A family-wise-error-corrected threshold of *P* < 0.05 was applied to all the contrasts discussed in this paper.

### fMRI model comparison

We compared three fMRI model’s ability to explain activity in the ventral striatum using two common methods. One, we calculated the AIC for the categorical, combined-RPE, and separated-RPE models for each voxel within a bilateral mask of the nucleus accumbens. Two, we estimated the models using SPM’s Bayesian estimation algorithm and compared them using Bayesian model selection.

### Acquisition and preprocessing of pupil diameter data

Pupil diameter was measured by an EyeLink 1000 eye tracker (SR Research) each 2 ms. Pupil diameter for each session was processed by first eliminating 100 ms around each blink. If the remaining pupil data for that session spanned less than than 80% of the session’s time, then all data for that session was eliminated from the analysis. In total, we analyzed data from 190 sessions and 73 participants (36 from the story condition and 37 from the abstract condition). Data from each included session were z-scored, smoothed with a median filter (window size 50) and a Gaussian filter (SD 10), and interpolated with a third-order spline interpolator, using the SciPy Python library [85].

The mean pupil diameter during feedback, first-stage choice, and interval between the first-stage and second-stage choices was obtained by taking pupil diameter values every 2 ms within each of these intervals for each trial and averaging them. These data were then analyzed with a hierarchical Bayesian regression using the brms package for R [86]. The feedback model included the factors reward, transition, and their interaction for each trial as random effects for each participant, as well as condition, reward, transition, and all their interactions as fixed effects across all participants. As a brms formula, it can be written as:

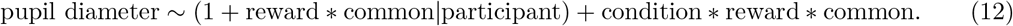

The first-stage model only had a fixed effect for the condition because the reward/no reward outcome was not yet revealed at that stage.

## Data availability

The behavioural and eye-tracking data can be found on https://github.com/carolfs/fmri_magic_carpet and the fMRI images can be found on [link to be determined].

## Code availability

The code used to run the task and the analyses can be found on https://github.com/carolfs/fmri_magic_carpet

## Acknowledgements

We would like to thank Giuseppe M Parente for the illustrations used in the experimental tasks, Karl Treiber and Elena Silingardi for helping with the fMRI data collection, Susanna Gobbi for helping with the fMRI preprocessing and analysis as well as reviewing our calculations, and ND Daw, P Dayan, M Grueschow, A Konovalov, I Krajbich, and S Nebe for helpful comments on early drafts of this manuscript. Our acknowledgement of their feedback does not imply that these individuals fully agree with our conclusions or opinions in this paper. This work was supported by the CAPES Foundation (Grant 88881.119317/2016-01) and the European Union’s Seventh Framework programme for research, technological development and demonstration under grant agreement no 607310 (Nudge-it). The funders had no role in study design, data collection and analysis, decision to publish or preparation of the manuscript.

## Author contributions

CFS and TAH conceived the project. All authors designed the experiments. CFS and GL collected and analyzed the data with input from ME and TAH. CFS and TAH wrote the first draft of the manuscript. All authors revised the manuscript for submission.

## Competing interests

The authors declare no competing interests.

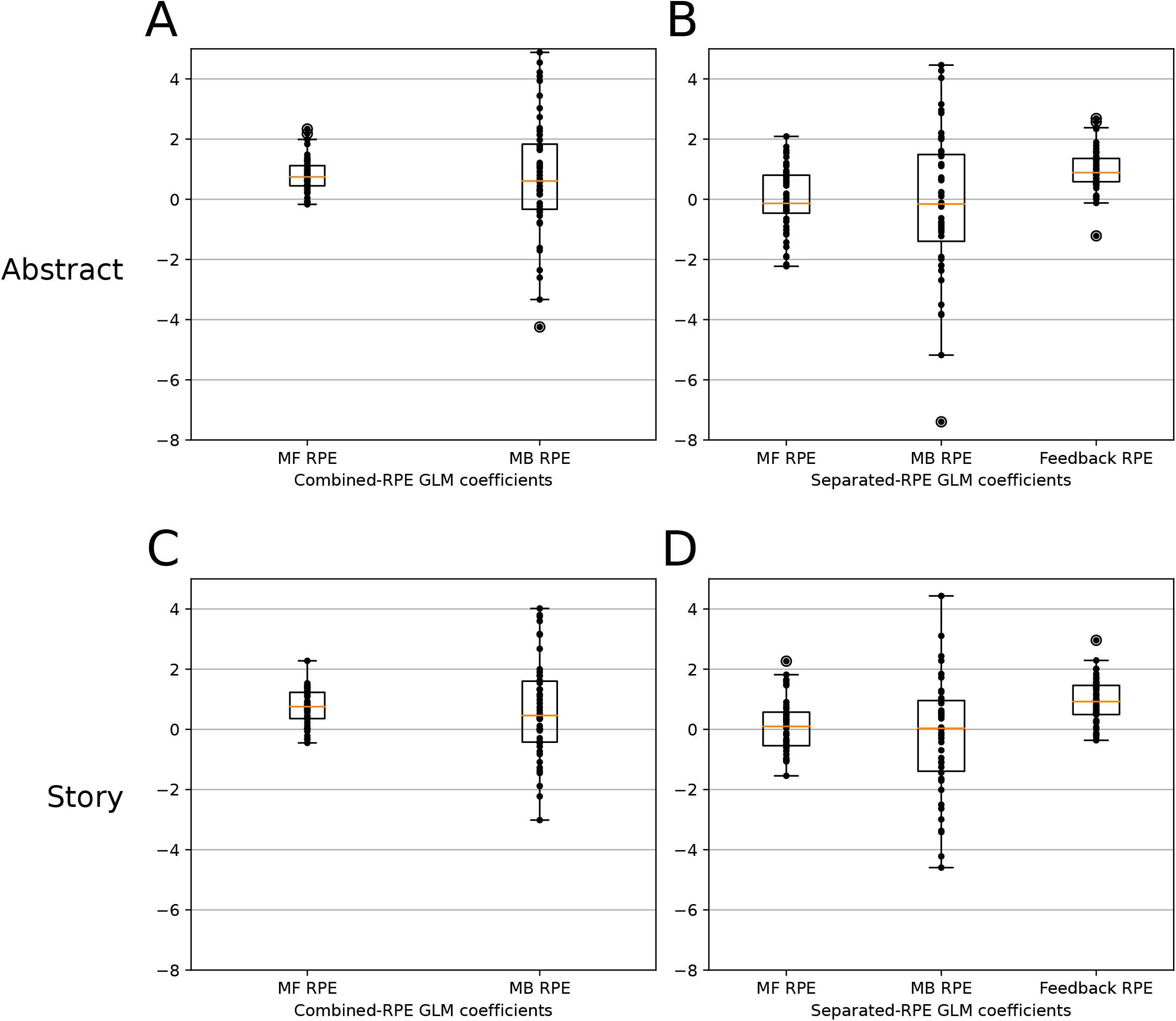

## Notes

https://github.com/carolfs/fmri_magic_carpet

### Competing Interest Statement

The authors have declared no competing interest.

