## Supplemental Material for "A new take on model-based and model-free influences on mental effort and striatal prediction errors"

3rd November 2022

#### Hybrid RL model parameters

We fit the hybrid reinforcement learning model [1] to our data from both instruction conditions. Descriptive statistics for the parameters are reported in Table S1. One parameter of interest in this model is the model-based weight $w$, which measures the balance between model-free and model-based influences on decisions taken at the first stage, under the critical assumption that those are the only two strategies a participant might use. A value of $w = 0$ indicates a purely model-free strategy, a value of $w = 1$ indicates a purely model-based strategy, and a value $0 < w < 1$ indicates a hybrid strategy. Consistent with the logistic regression results, the model-based weight is higher for the story than abstract instruction condition with 0.89 probability.

For the abstract condition, PSIS-LOO IC $\pm$ SE: $6791.7 \pm 94.5$ for the hybrid reinforcement learning (RL) model and $6600.8 \pm 91.3$ for the logistic regression; for the story condition, PSIS-LOO IC $\pm$ SE: $8455.5 \pm 149.6$ for the hybrid RL model and $6829.2 \pm 81.0$ for the logistic regression.

| Parameter | Abstract | Story |
|---|---|---|
| $\alpha_1$ | 0.14 $[0.03, 0.32]$ | 0.08 $[0.00, 0.28]$ |
| $\alpha_2$ | 0.51 $[0.36, 0.65]$ | 0.88 $[0.78, 0.96]$ |
| $\lambda$ | 0.97 $[0.57, 1.00]$ | 0.89 $[0.15, 1.00]$ |
| $\beta_1$ | 6.57 $[5.45, 7.78]$ | 6.72 $[5.43, 8.05]$ |
| $\beta_2$ | 3.72 $[3.19, 4.28]$ | 2.60 $[2.23, 3.01]$ |
| $w$ | 0.60 $[0.46, 0.75]$ | 0.73 $[0.58, 0.86]$ |
| $p$ | 0.14 $[0.11, 0.18]$ | 0.04 $[0.02, 0.07]$ |

Table S1: Hybrid RL model parameters per instruction condition (median and 95% High-Posterior-Density Interval). These parameters were estimated using a hierarchical Bayesian model with a condition regressor on the median hyperparameters, and the values listed are derived from the resulting distributions for each instruction condition.

#### Second-stage reaction time analysis confirms that participants in the story condition used a more correct model-based strategy than participants in the abstract condition

Previous findings suggest that participants with a more correct model-based strategy are faster to respond at the second stage after a common transition compared to a rare transition [2, 3]. We looked for this effect in our behavioural data to further support our conclusion that participants in the story condition used a more correct model-based strategy than participants in the abstract condition. We ran a linear regression model where log-transformed second-stage reaction times ($\log(\mathrm{RT}_2)$) were a function of transition (coded as common = 1 or rare = 0), condition (coded as story = 1 or abstract = 0), and their interaction, with participant as a

| Effect | Estimate | Std.Error | DF | $t$ value | $P$ value |
|---|---|---|---|---|---|
| intercept | 0.03963 | 0.02268 | 92.30488 | 1.747 | 0.0839 |
| story | 0.06007 | 0.03243 | 92.40279 | 1.852 | 0.06719 |
| common | -0.17974 | 0.02253 | 91.89436 | -7.977 | $4.5 \times 10^{-12}$ |
| story:common | -0.09276 | 0.03222 | 91.97958 | -2.879 | 0.00496 |

Table S2: Analysis of log-transformed second-stage reaction times as a function of transition (coded as common = 1 or rare = 0), condition (coded as story = 1 or abstract = 0), and their interaction, with participant as a random effect on the intercept and transition. These results suggest that participants in the story condition adopted a more correct model-based strategy than participants in the abstract condition.

random effect on the intercept and transition:

$$\log(\mathrm{RT}_2) \sim (1 + \mathrm{common}|\mathrm{participant}) + \mathrm{story} * \mathrm{common}. \quad (1)$$

This model was then fitted to the behavioural data using the `lmer` function of the R packages lme4 [4] and lmerTest [5]. The results are as shown in Table S2. They suggest that participants were faster to respond after a common transition compared to a rare transition, and this effect was larger for the story condition compared to the abstract condition, which supports our conclusion. Second-stage reaction times are plotted in Figure S1.

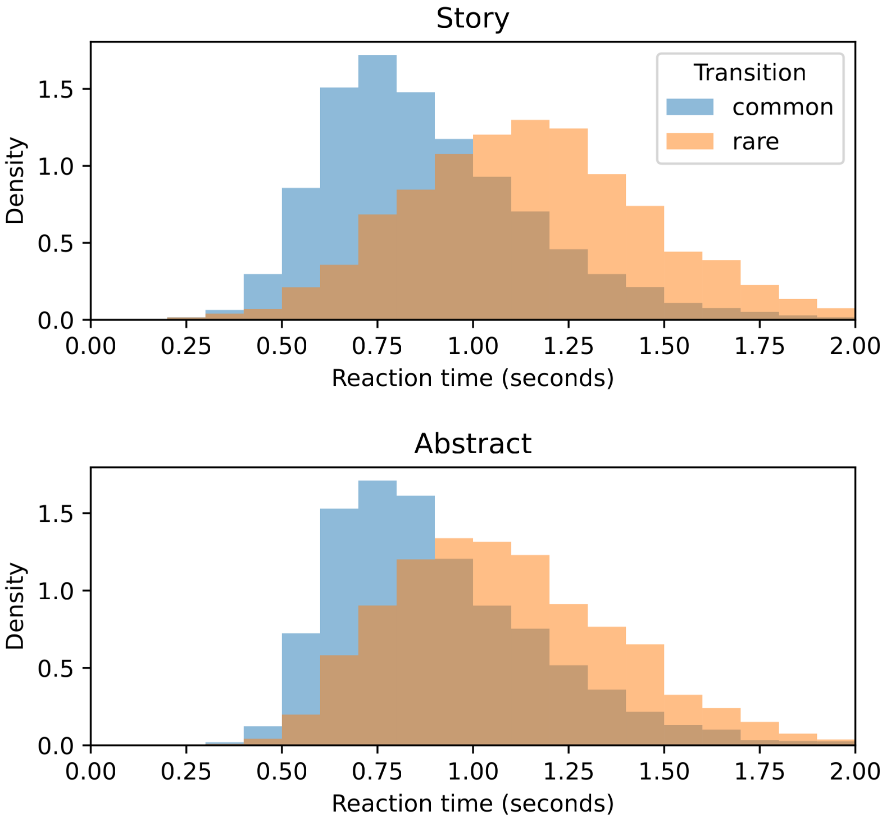


Figure S1: Histograms of second-stage reaction times for the story and abstract conditions, after common versus rare transitions. Participants are faster to respond after a common transition compared to a rare transition, especially in the story condition. This reaction time difference is an evidence of correct model-based behaviour.

#### Updating and decision processes after feedback

Consistent with the theory of reward learning in general, the previous results from Konovalov and Krajbich [6] indicate that mental model updates occur after feedback rather than at the first stage decision, where participants actually use a given strategy. Those authors examined how the number of visual fixations at the first stage differed between participants classified as more model-based or more model-free. They observed that more model-based participants made fewer fixations at the first stage, as if they had already made their choices right after feedback without waiting for the first-stage choices to be displayed again, This is likely because first-stage choices are always the same and thus known in advance. In our experiment, participants had 2–7 seconds between feedback and the next trial, while there was no delay between trials in Konovalov and Krajbich [6], making it even more likely that our participants decided before first-stage onsets. In support of this hypothesis, we found that the median reaction time at the first stage is not significantly correlated with the model-based weight (Spearman's $\rho = 0.09$, $P = 0.37$; the model-based weight for each participant was estimated by the median of the posterior distribution) or with the transition-by-reward coefficient in the logistic regression analysis of consecutive trial pairs (Spearman's $\rho = -0.13$, $P = 0.21$; the coefficient for each participant was estimated by the median of the posterior distribution).

#### Functional MRI results

We did not detect any significant differences in BOLD activity between the two conditions at the first stage, which is consistent with the hypothesis that first-stage decisions were made in advance, right after feedback. There were significant differences between conditions at the second stage (Table S3. We had no a priori predictions about group differences at the second stage, but a Neurosynth [7] analysis indicates that they are associated with the anticipation and execution of motor responses (the top 10 terms are: motor, motor cortex, primary motor, primary, hand, supplementary motor, movement, supplementary, anticipation, movements).

| Harvard-Oxford Probabilistic Atlas labels | Peak MNI coordinates (mm) | | | Number of Voxels |
|---|---|---|---|---|
| | x | y | z | |
| **Second stage** | | | | |
| 35% Postcentral Gyrus, 13% Precentral Gyrus | -39 | -27 | 69.6 | 106 |
| 34% Precentral Gyrus, 14% Postcentral Gyrus | 36 | -24 | 69.6 | 26 |
| 35% Right Accumbens, 32% Right Caudate, 2% Subcallosal Cortex | 6 | 12 | -2.4 | 16 |
| 46% Paracingulate Gyrus, 9% Juxtapositional Lobule Cortex (formerly Supplementary Motor Cortex), 4% Cingulate Gyrus, anterior division | 0 | 9 | 48 | 9 |
| **Feedback** | | | | |
| 22% Frontal Orbital Cortex, 22% Inferior Frontal Gyrus, pars triangularis, 14% Frontal Operculum Cortex, 3% Inferior Frontal Gyrus, pars opercularis | 51 | 21 | -6 | 211 |
| 64% Frontal Pole, 9% Frontal Orbital Cortex, 6% Inferior Frontal Gyrus, pars triangularis | 51 | 39 | -13.2 | 22 |
| 9% Frontal Orbital Cortex, 8% Insular Cortex, 6% Temporal Pole | -33 | 12 | -20.4 | 8 |
| 44% Right Caudate | 6 | 9 | 1.2 | 2 |

Table S3: This table reports voxel clusters that exhibited higher BOLD activity in participants in the abstract instruction condition ($N = 48$) compared to the story instruction condition ($N = 45$) during the second stage and feedback. No differences in activation were above threshold during the first stage. Peaks whose $x$ coordinate is negative/positive are located in the left/right hemisphere respectively. Labels correspond to structures to which each cluster peak belongs. Each voxel measures $3 \times 3 \times 3\text{mm}^2$. (Threshold: family-wise-error-corrected, $P < 0.05$.)

| Harvard-Oxford Probabilistic Atlas labels | Peak MNI coordinates (mm) | | | Number of Voxels |
|---|---|---|---|---|
| | x | y | z | |
| **Common > rare** | | | | |
| 42% Parietal Operculum Cortex, 34% Planum Temporale, 5% Supramarginal Gyrus, posterior division, 5% Superior Temporal Gyrus, posterior division, 2% Supramarginal Gyrus, anterior division | -57 | -36 | 19.2 | 1003 |
| 32% Planum Polare, 32% Central Opercular Cortex, 7% Precentral Gyrus, 5% Superior Temporal Gyrus, anterior division, 2% Planum Temporale, 2% Heschl's Gyrus (includes H1 and H2) | 60 | 0 | 4.8 | 894 |
| 22% Frontal Pole, 1% Superior Frontal Gyrus | -6 | 60 | 37.2 | 152 |
| 71% Temporal Pole | -54 | 9 | -24 | 146 |
| 25% Frontal Orbital Cortex, 23% Inferior Frontal Gyrus, pars triangularis, 20% Frontal Pole | -48 | 33 | -6 | 144 |
| 42% Frontal Medial Cortex, 30% Frontal Pole | 0 | 54 | -16.8 | 120 |
| 43% Precentral Gyrus, 25% Postcentral Gyrus | -45 | -12 | 33.6 | 73 |
| Cerebellum | 27 | -81 | -34.8 | 13 |
| 42% Postcentral Gyrus, 22% Superior Parietal Lobule, 1% Precentral Gyrus | 24 | -39 | 62.4 | 5 |
| 44% Postcentral Gyrus, 25% Supramarginal Gyrus, anterior division | 60 | -18 | 37.2 | 3 |
| 21% Superior Parietal Lobule, 3% Postcentral Gyrus | 21 | -48 | 76.8 | 3 |
| 12% Frontal Pole | -9 | 48 | 51.6 | 2 |
| **Rare > common** | | | | |
| 19% Angular Gyrus, 18% Lateral Occipital Cortex, superior division, 11% Superior Parietal Lobule, 6% Supramarginal Gyrus, posterior division | -33 | -57 | 37.2 | 4408 |
| 51% Paracingulate Gyrus, 3% Superior Frontal Gyrus, 2% Juxtapositional Lobule Cortex (formerly Supplementary Motor Cortex), 1% Cingulate Gyrus, anterior division | 0 | 15 | 48 | 292 |
| 51% Middle Frontal Gyrus, 18% Frontal Pole, 7% Inferior Frontal Gyrus, pars triangularis | -45 | 33 | 22.8 | 248 |
| 37% Middle Frontal Gyrus, 14% Precentral Gyrus, 13% Inferior Frontal Gyrus, pars opercularis | 39 | 12 | 30 | 164 |
| 45% Frontal Orbital Cortex, 21% Insular Cortex | -30 | 24 | -6 | 115 |
| 63% Frontal Orbital Cortex, 5% Insular Cortex | 33 | 24 | -9.6 | 107 |

Table S4: This table reports voxel clusters that exhibited higher BOLD activity after common transitions compared to rare transitions and vice versa in the abstract instruction condition ($N = 48$). Peaks whose $x$ coordinate is negative/positive are located in the left/right hemisphere respectively. Labels correspond to structures to which each cluster peak belongs. Each voxel measures $3 \times 3 \times 3\text{mm}^2$. (Threshold: family-wise-error-corrected, $P < 0.05$.)

### Replicating hybrid RL model prediction error correlations with ventral striatal activity

To test whether our data from participants in the abstract condition were consistent with past reports, we attempted to replicate the central fMRI findings in those studies. The results reported by Daw et al. [1] have already been replicated several times [2, 8, 9]. In order to better match the methodology employed in previous studies, we refit the hybrid reinforcement learning model to our data using a mixed-effects approach, where only the model-based weights were allowed to vary between participants and all the other parameters were constrained to be the same for all participants[1] (see Table S5). Using these parameters, we generated model-free and model-based reward prediction errors (RPEs) for each trial at the second stage choice and feedback events. We then ran the combined-RPE fMRI model used in previous publications and tested for BOLD activity that

[1]This approach was advocated by Daw et al. [1] because: "in these sorts of algorithms, noise and variation in parameter estimates from subject to subject results, effectively, in a rescaling of regressors between subjects, which suppresses the significance of neural effects in a subsequent second-level fMRI analysis, producing poor results."

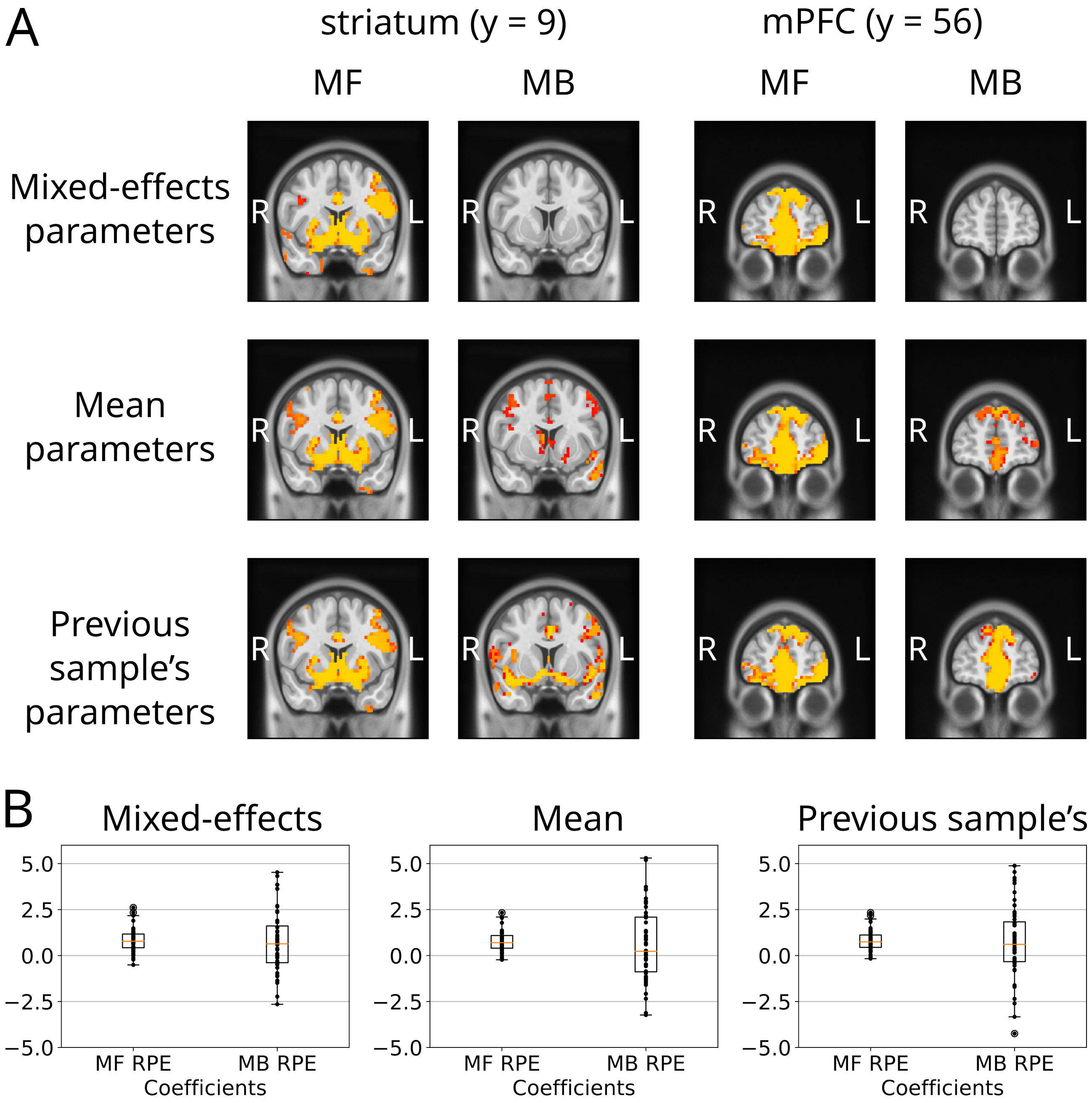


Figure S2: fMRI results for the combined-RPE GLM using different parameter sets. A) Each row shows the correlation between BOLD activity in the striatum and mPFC and model-free (MF) or model-based (MB) RPEs for participants in the abstract condition. The RPE regressors in each row were computed using the different sets of hybrid RL model parameters listed in Table S5. The results shown in the third row are based on the parameters fit to the choices made by participants in a previous study [1]. B) Mean coefficients for participants in the abstract condition within the nucleus accumbens. Each black dot represents the mean coefficient from a single participant. The box and whisker plots show the distribution across the entire sample. The box extends from the first quartile to the third quartile of the distribution, with a line at the median. The whiskers extend from the box by 1.5 times the inter-quartile range.

| Sample | Method | $\alpha_1$ | $\alpha_2$ | $\lambda$ | $\beta_1$ | $\beta_2$ | $p$ |
|---|---|---|---|---|---|---|---|
| Current abstract | mixed effects | 0.16 | 0.50 | 0.44 | 6.13 | 3.01 | 0.15 |
| Current story | mixed effects | 0.01 | 0.86 | 0.84 | 8.53 | 2.00 | 0.04 |
| Current abstract | mean estimates | 0.48 | 0.57 | 0.62 | 6.72 | 3.81 | 0.24 |
| Current story | mean estimates | 0.55 | 0.78 | 0.51 | 6.33 | 2.68 | 0.07 |
| Previous [1] | mixed effects | 0.70 | 0.40 | 0.63 | 4.23 | 2.95 | 0.17 |

Table S5: This table reports the hybrid reinforcement learning model parameter estimates we tested in our fMRI analyses. The first column identifies the source of the choice data the model was fit to in order to obtain the parameter estimates. The parameter estimates in the first two rows came from fitting the choice data from participants in our abstract instructions condition using two different methodologies. In the fMRI replication attempt, we used only the data from participants in the abstract condition to better align with the procedures used in previous fMRI studies with the two-stage task. The parameters in the third row were fit to the choice data from a previous sample of participants reported in Daw et al. [1]. The second column indicates the method used to obtain the estimates (either by fitting a mixed-effects model to the complete data set or by fitting the model to each participant separately and subsequently taking the mean of those individual fits). The subsequent six columns list the estimates for the model's parameters, which were used to calculate reward prediction error regressors for the fMRI analyses of all participants in the abstract instructions condition.

significantly correlated with these reward prediction errors after correcting for multiple comparisons ($p < 0.05$, whole-brain corrected). Consistent with previous reports [1, 2, 8, 9], the model-free reward prediction error was significantly correlated with activity in the bilateral ventral striatum and areas of the medial prefrontal cortex (mPFC); however, in contrast, the model-based reward prediction error did not show significant associations with BOLD signals in any brain region in our sample of participants (first row of Figure S2). Several previous studies [1, 2, 8, 9] have found significant associations between model-based prediction errors and BOLD signals in the striatum and mPFC. Therefore, we investigated the possible reasons why this result did not replicate in our data from the abstract instruction condition. We ran exploratory analyses on the data from the story instruction group and found similar results. However, we stick to our a priori plan of trying to replicate previously reported findings within the better-matching abstract instruction group.

In the course of this investigation, we found that the hybrid RL parameters used to calculate the RPEs had a strong impact on the association between RPEs and striatal BOLD activity. We noted that the best-fitting parameters we obtained by fitting the mixed-effects hybrid model to our data were significantly different from the parameters obtained by Daw et al. [1]. Furthermore, other papers that replicated the fMRI results in Daw et al. [1] used the mean parameter values across all participants to calculate the reward prediction errors instead of fitting a mixed-effects model to the data (e.g. [8]). Therefore, we ran the fMRI analysis twice more, using either the mean parameter values from our participants or the parameter values reported in Daw et al. [1] to calculate the RPEs (see Figure S3 for histograms of the RPEs calculated using the latter parameters). These two analyses both revealed a correlation between the model-based reward prediction error and activity in the ventral striatum and the medial prefrontal cortex, consistent with previous papers. Interestingly, using the parameters previously reported by Daw et al. [1] yielded the largest number of voxels in mPFC and ventral striatum significantly correlating with model-based prediction errors even though those parameters did not best explain our participants choice data (Figure S2). The discovery that the fMRI results for the model-based prediction error regressor depended heavily on the hybrid reinforcement learning model parameters used to compute the prediction error prompted us to examine the prediction error regressors within the combined-RPE GLM in more detail. Even though the combined-RPE GLM did not best explain the BOLD activity in the ventral striatum, we thought it important to examine the combined-RPE GLM to better understand and interpret the previously reported results derived from that model [1, 2, 8, 9].

### Averaging RPE effects over second stage and feedback events may produce misleading results

The combined-reward-prediction-error (combined-RPE) fMRI model is restricted to quantifying how BOLD activity relates to reward prediction errors at the second-stage and feedback events with a single regression coefficient. In other words, the regression coefficients for model-based and model-free RPEs represent the average across both time points. Therefore, none of the results about prediction errors in the previous studies using this fMRI model specification (i.e., all of them) apply exclusively to the second stage, which is the only moment in a trial when the brain would calculate distinct model-free and model-based RPEs according to the hybrid reinforcement learning algorithm. Concretely, the combined-RPE fMRI model contains the following regressors (excluding motion and other confound regressors).

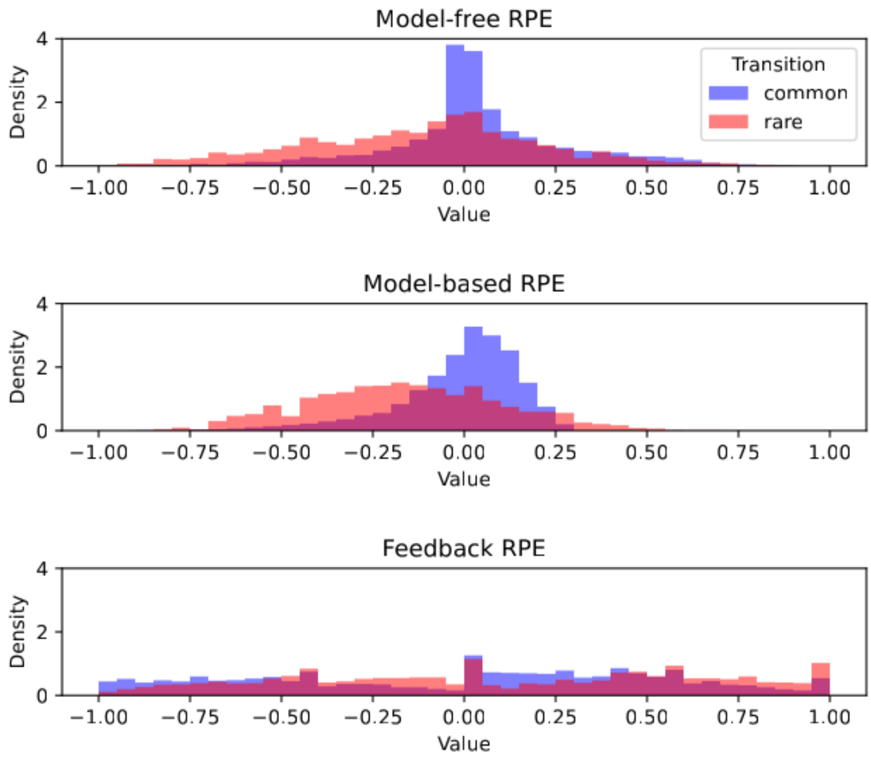


Figure S3: Histograms of the three types of RPEs in the two-stage task—model-free, model-based, and feedback—after a common or a rare transition, calculated using the parameters from the original study [1]

.

1. a binary regressor for first-stage choice onsets, which were also modulated by:
   (a) the value of the chosen first-stage action
   (b) the partial derivative of the chosen value with respect to $w$
2. a binary regressor combining second stage and feedback events, which were also modulated by:
   (a) model-free RPEs ($RPE_{MF}$) combined across the second stage and feedback.
   (b) the difference between the model-based and model-free RPEs ($RPE_{\Delta MB}$) combined across the second stage and feedback.
3. a binary regressor for feedback onsets without any parametric modulators

According to the authors of the original publication, the difference between the RPEs was employed instead of just the model-based RPE itself in order "to reduce the correlation between the regressors of interest, and also because it encompassed the test of the null hypothesis that RPE signaling in striatum was purely model-free" [1]. The subtraction step used to create $RPE_{\Delta MB}$ means that instead of the positive correlation between $RPE_{MF}$ and $RPE_{MB}$ at the second stage, we now have a negative correlation between $RPE_{MF}$ and $RPE_{\Delta MB}$ at this time point, with the strengths of these correlations depending on the exact hybrid model parameters. We note that in addition to the steps the authors took to reduce correlations between the RPE regressors, by default the SPM software (versions 5 and 8) used in all the previously published fMRI studies of the two-stage task mean-centers and orthogonalizes the parametric modulators in the order they are entered, before and after convolution with the hemodynamic response function. Despite this orthogonalization process, the specific combination of regressors in the combined-RPE GLM results in a negative correlation between $RPE_{MF}$ and $RPE_{\Delta MB}$ at the critical second-stage choice event (we provide a set of simulations demonstrating this effect here: `https://colab.research.google.com/drive/1c_-firW7x9NfRkNyzUKpsFPV3TLahOV7#scrollTo=RTR9ePJs29ML`). The bottom line is that, in the context of this combined-RPE GLM, there is a negative correlation between the $RPE_{MF}$ and $RPE_{\Delta MB}$ at the second stage with or without the orthogonalization step.

The negative correlation between the two RPE regressors at stage two together with the fact that RPE correlations with BOLD activity at the second stage and feedback are quantified by the same regression coefficient biases the apparent correlation between BOLD activity and $RPE_{\Delta MB}$. The type and strength of the bias depend on the size and direction of the RPE correlations at the second-stage and feedback.

Although the hybrid algorithm does not distinguish between $RPE_{MF}$ at the second stage and feedback, the brain may respond differently than that algorithm predicts. Our current data clearly indicate that the correlations between $RPE_{MF}$ and BOLD activity differ at the second stage versus feedback (see Figure S5) both for the abstract and the story conditions. The separated-RPE GLM shows that the $RPE_{MF}$ does not exhibit significant positive correlations with brain activity at the second stage (Figures S4 and S5)[2], but BOLD activity at feedback does positively correlate with reward prediction errors (again, $RPE_{MF}$ and $RPE_{MB}$ are the same at feedback). Moreover, for every participant in the abstract condition, the mean effect of the feedback RPE on BOLD activity within the nucleus accumbens was higher than the mean effect of the second-stage model-free RPE, and this difference was statistically significant (two-sided Wilcoxon signed rank test, median difference: 0.906, 95% CI $[0.618, 1.281]$, $V = 1055$, $P < 0.001$).

Thus, the results for the model-free regressor when the combined-RPE is fit to data from our sample—and potentially other papers using the same GLM—are actually driven by how well this predictor correlates with brain activity at feedback. Critically, the model-free and model-based prediction errors are identical at feedback, meaning that this result provides no evidence for the separate reward prediction error signals we would expect if the brain employs both learning algorithms during the two-stage task.

The apparent correlations with $RPE_{\Delta MB}$ at the second-stage in the combined-RPE GLM are the result of the biased estimates this regression specification generates for its coefficient. As noted above, the estimated coefficient for $RPE_{\Delta MB}$ depends on its correlation with $RPE_{MF}$ and the difference in how well $RPE_{MF}$ correlates with BOLD activity at the second stage versus feedback events. The fact that the degree of bias depends on the strength of the correlation between $RPE_{MF}$ and $RPE_{\Delta MB}$ explains the different results we find for the $RPE_{\Delta MB}$ regressor when computing it using different hybrid RL model parameters (Figure S2). The parameters we obtain when fitting our participants' choices using a mixed-effects approach generate a weaker correlation between $RPE_{MF}$ and $RPE_{\Delta MB}$ than taking the means across individual fits to each participant in our sample or using the mixed-effects fit reported in the original paper [1] (Figure S6, Table S5). The lower negative correlation between $RPE_{MF}$ and $RPE_{\Delta MB}$ from the mixed-effects fits to our data results in less positively biased coefficients for $RPE_{\Delta MB}$ in the combined-RPE GLM that do not significantly differ from zero. In contrast, the stronger biases generated by the other hybrid RL parameter combinations led to results that appear to be statistically significant, but are in fact spurious artifacts caused by the combined-RPE GLM specification. This was confirmed by deleting $RPE_{MF}$ from the separated-RPE GLM and re-fitting the model to our data; no correlation was found between brain activity in the nucleus accumbens and $RPE_{\Delta MB}$.

[2]We used mean-centering and orthogonalization of the parametric modulators in the separated-RPE GLM analysis for consistency with previous studies, but omitting these transformations of the regressors only changed which areas were negatively correlated with the model-free RPE.

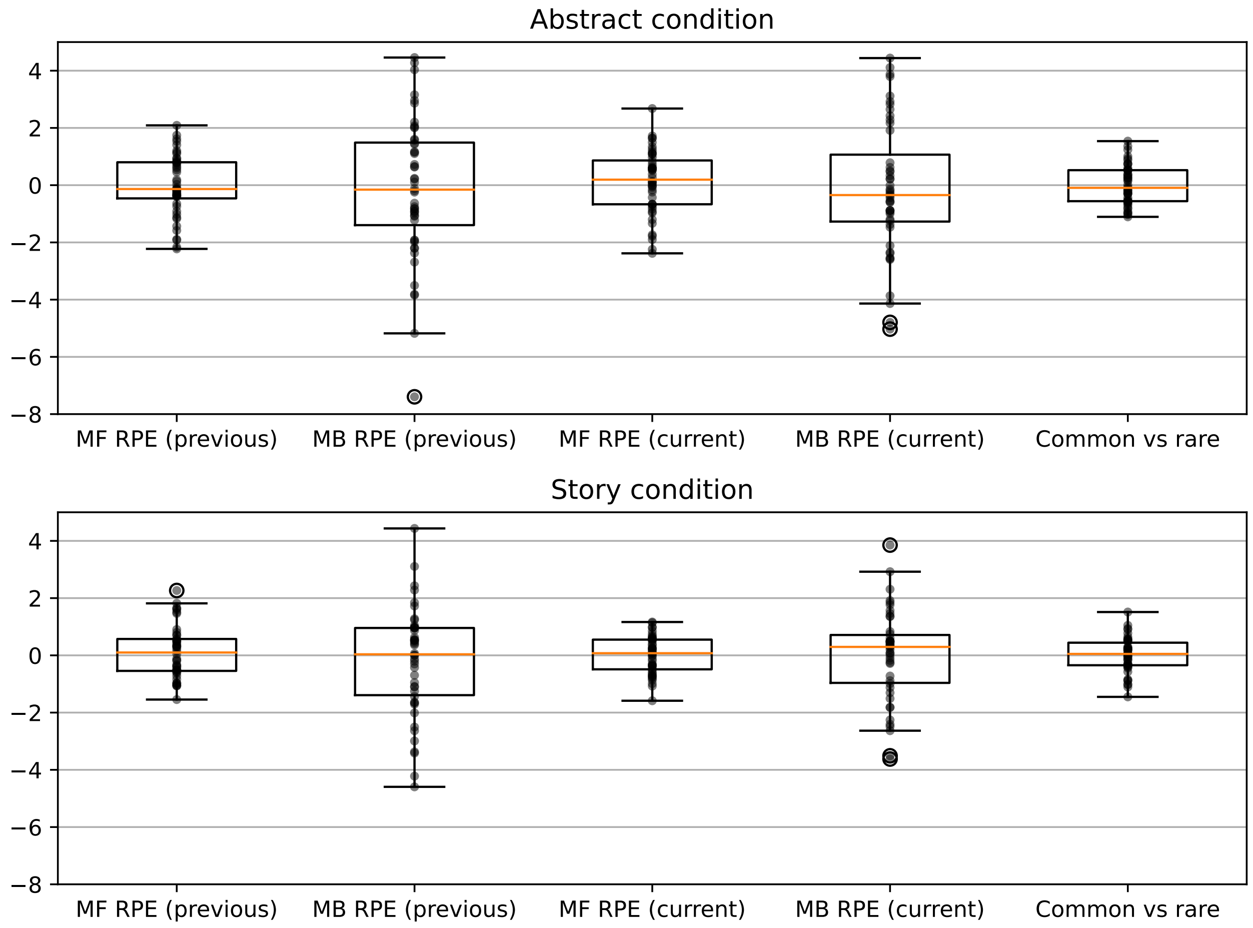


Figure S4: Mean estimated coefficients from the categorical (common vs. rare) and separated-RPE (second-stage model-free and model-based RPEs) GLMs within the nucleus accumbens for the abstract ($N = 48$) and story ($N = 46$) conditions. All of these regressors were included in the models to explain BOLD activity at the second stage but not at feedback. The model-free and model-based RPEs were calculated using the previous study's parameters [1] in one analysis (first two data sets of each subplot) and the current study's mixed-effect parameters in another analysis (next two data sets of each subplot). In either one-sample one-sided t-tests or Wilcoxon signed-rank tests, none of the means of these coefficient sets were found to be significantly greater than zero, with the minimum uncorrected $P$-value being 0.195. Each black dot represents the coefficient from a single participant. The box and whisker plots show the distribution across the entire sample. The box extends from the first quartile to the third quartile of the distribution, with a line at the median. The whiskers extend from the box by 1.5 times the inter-quartile range.

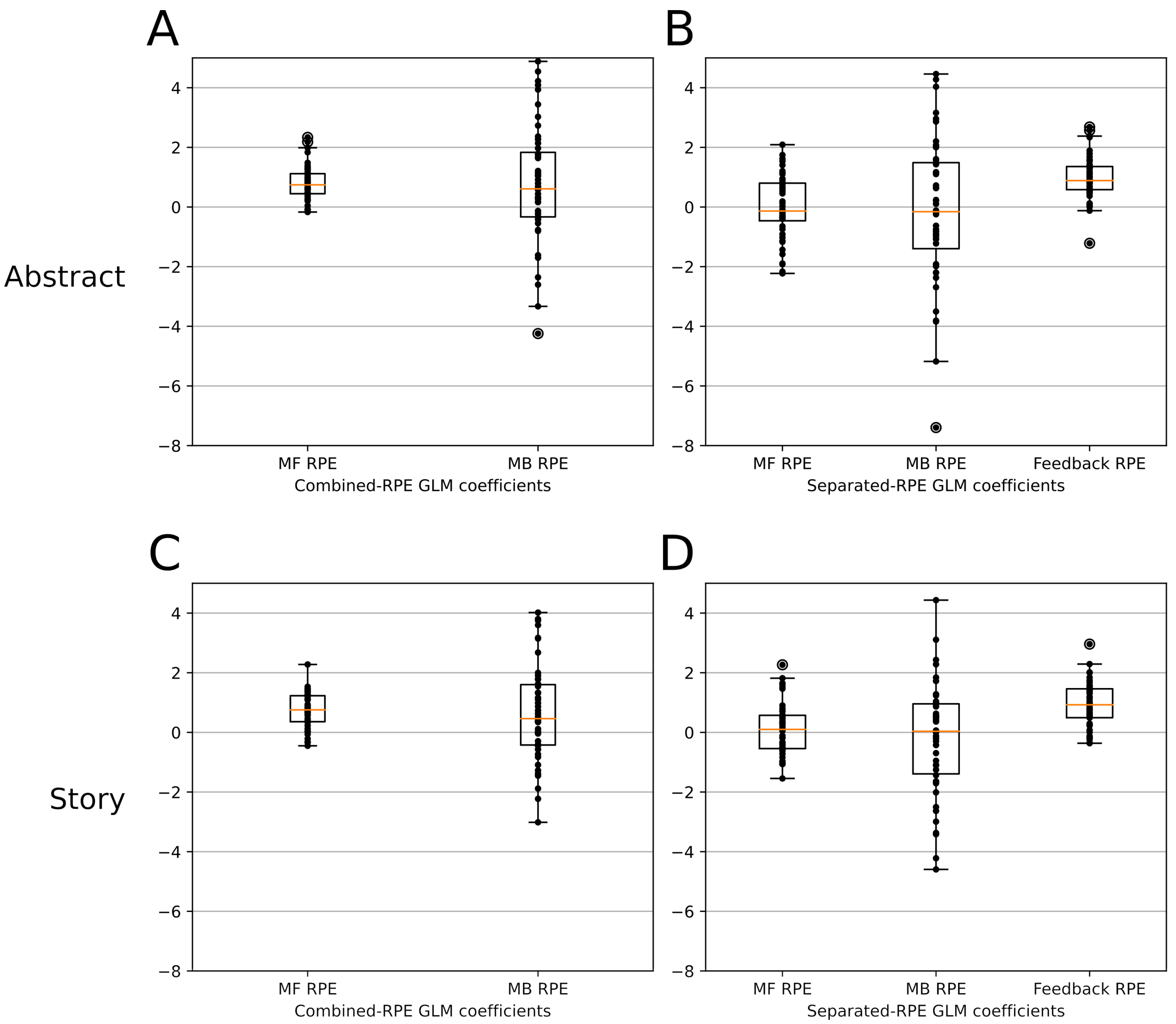


Figure S5: Mean estimated coefficients from the A, C) combined-RPE and B, D) separated-RPE GLMs within the nucleus accumbens for the abstract ($N = 48$) and story ($N = 46$) conditions. Each black dot represents the coefficient from a single participant. The box and whisker plots show the distribution across the entire sample. The box extends from the first quartile to the third quartile of the distribution, with a line at the median. The whiskers extend from the box by 1.5 times the inter-quartile range. These coefficients were obtained using hybrid parameter estimates from the previous sample [1].

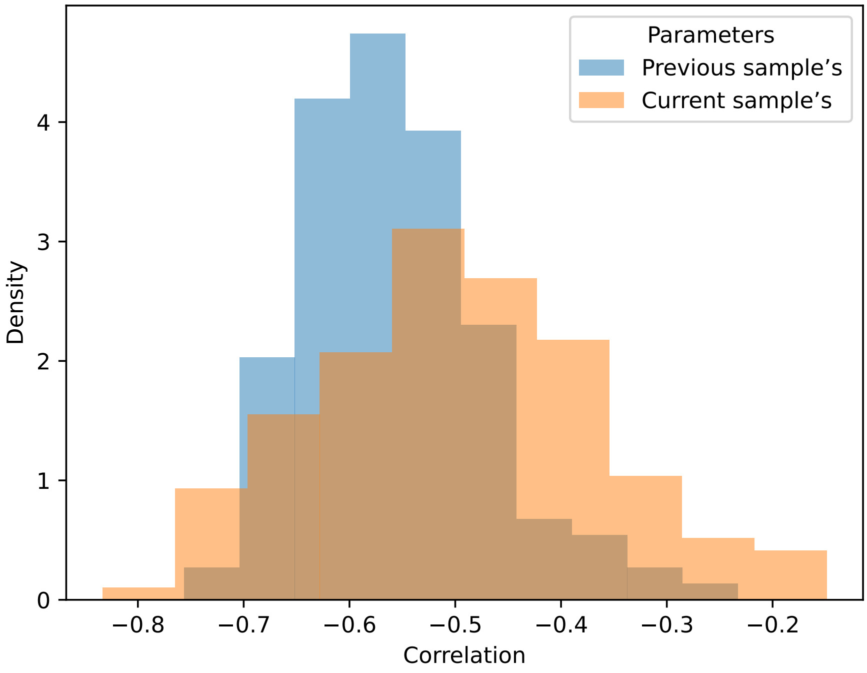


Figure S6: The magnitude of the correlation between model-free and model-based reward prediction errors regressors depends on the fitted hybrid reward learning model parameter estimates. The histograms show magnitude of the negative correlation between the parametric regressors for $RPE_{MF}$ and $RPE_{\Delta MB}$ (orthogonalized with respect to the combined-RPE GLM $RPE_{MF}$ regressor) at stage two for each of the three task runs completed by participants in the abstract instruction group. (To obtain the regressor for the $RPE_{\Delta MB}$ only at stage two, we "regressed out" the feedback onset regressor from the combined $RPE_{\Delta MB}$ regressor.) The orange bars show the correlations when the regressors are computed using the best-fitting parameters from a mixed-effects fit to the data from our sample. The blue bars show the correlations when the regressors are computed for the same participants using the best-fitting parameters from a mixed-effects fit to the data from a previous sample of participants [1]. In both cases, the parametric regressors are generated from the choices made by the participants in our sample, and the hybrid RL parameters are used to compute the corresponding reward prediction errors. For 79% of participants, the mean correlation across the three task runs was more negative when using the previous sample's hybrid RL parameters than when using those from the current sample. In parallel, we observed apparently significant, but in fact spurious, correlations between the parametric modulators for $RPE_{MF}$ and $RPE_{\Delta MB}$ from the combined-RPE GLM and BOLD activity in the nucleus accumbens at stage two when using the previously reported hybrid RL parameters to generate the RPEs, but not when the RPEs were based on the current sample's parameters (see Figure S2).

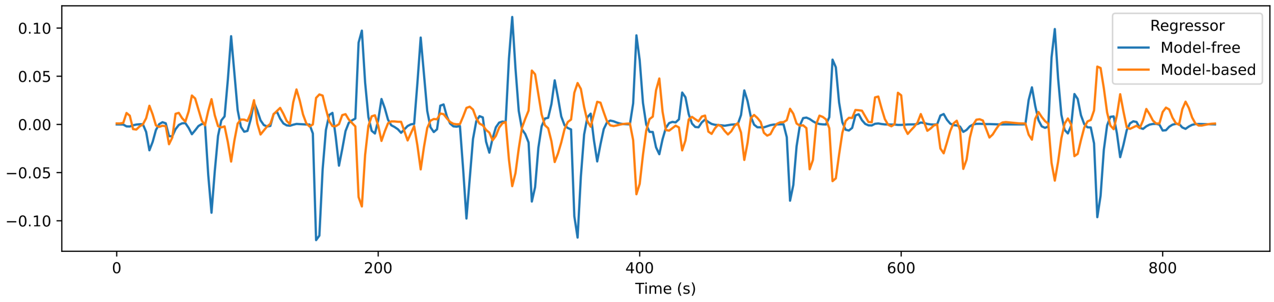


Figure S7: The model-free and orthogonalized model-based RPE regressors are negatively correlated at the second stage in the combined-RPE GLM. Here we plotted these regressors for run 3 of participant 25, which exhibits the most negative correlation in our data set ($\rho = -0.76$). Clearly, these regressors tend to make opposite predictions for BOLD activity and thus could largely cancel each other out if combined with optimized positive coefficients.

The equations below show how the separated and combined-RPE GLMs attempt to explain the fMRI data $y$ as a function of the reward prediction error signals at the second-stage decision and feedback events.

Equation 1 represents an abbreviated version of the separated-RPE GLM and includes the parametric regressors $RPE_{MF}$ and $RPE_{\Delta MB}$ at the second-stage decision (S2) and a parametric regressor for the RPE at feedback (FB), which is the same for the MF and MB algorithms (the equation below omits an intercept and separate binary onset regressors for the second stage and feedback for conciseness):

$$y = \beta_1 S2_{RPE_{MF}} + \beta_2 S2_{RPE_{\Delta MB}} + \beta_3 FB_{RPE} + \ldots \tag{2}$$

Equations 2-3 shows the combined-RPE GLM which restricts the regression coefficients for RPEs to be the same at second stage decision and feedback events.

$$y = \beta_{C1} S2_{RPE_{MF}} + \beta_{C2} S2_{RPE_{\Delta MB}} + \beta_{C1} FB_{RPE_{MF}} + \beta_{C2} 0 + \ldots \tag{3}$$

We list $\beta_{C1}$ in red to highlight the fact that it quantifies the relationship between RPEs and BOLD activity at two separate points in the task (S2 and FB). In contrast, $\beta_{C2}$ only quantifies the relationship between $RPE_{\Delta MB}$ and BOLD activity at stage two because $RPE_{\Delta MB}$ equals zero at feedback. Thus, we can drop the final term, $\beta_{C2}0$, and we are left with,

$$y = \beta_{C1} S2_{RPE_{MF}} + \beta_{C2} S2_{RPE_{\Delta MB}} + \beta_{C1} FB_{RPE_{MF}} + \ldots \tag{4}$$

Equations 1 and 3 will generate similar results only when the association between $RPE_{MF}$ and $y$ is constant at stage two (S2) and feedback (FB). However, this is not the case in our data, and this critical assumption was not explicitly tested in previous work [1, 2, 8, 9]. Fitting equation 1 (i.e., the separated-RPE GLM) to our data shows that the correlation between $y$ and $S2_{RPE_{MF}}$ is lower than the correlation between $y$ and $FB_{RPE}$ (Figure S5). When fitting equation 3 to the same data, $\beta_{C1}$ must explain both correlations. Thus it would need to take on a compromise value between 0 and the value of the correlation at feedback, resulting in substantial residual error at both time points. However, if $S2_{RPE_{MF}}$ and $S2_{RPE_{\Delta MB}}$ are correlated, then $\beta_{C2}$ can offset $\beta_{C1}$ at S2 because $\beta_{C2}$ only influences the model fit at S2 and the value of $y$ at stage two is predicted by $\beta_{C1} S2_{RPE_{MF}} + \beta_{C2} S2_{RPE_{\Delta MB}}$. When the correlation between $S2_{RPE_{MF}}$ and $S2_{RPE_{\Delta MB}}$ is negative, $\beta_{C2}$ takes on a positive value to cancel out $\beta_{C1}$ at stage two (Figure S7). In our data, the combined orthogonalized $S2_{RPE_{\Delta MB}}$ is negatively correlated with the $S2_{RPE_{MF}}$ regressor for all runs and all participants when using the hybrid parameters from the previous sample reported in Daw et al. [1] (mean correlation: $-0.56$, SD: 0.09, range: $[-0.76, -0.23]$; Figure S6). Thus, the value of $\beta_{C1}$ is positive because it must explain the association between $y$ and $FB_{RPE}$ and the value of $\beta_{C2}$ is spuriously positive in order to cancel out $\beta_{C1}$ at the second stage.

### Pupil results

#### Additional details on the pupil diameter results at feedback

Table S6 below lists all the group-level effects in our analysis of pupil diameter at feedback.

| Predictors | Mean estimate | 95% CI |
|---|---|---|
| intercept | 0.43 | $[+0.36, +0.49]$ |
| condition | $-0.09$ | $[-0.15, -0.02]$ |
| reward | 0.05 | $[+0.03, +0.08]$ |
| transition | $-0.07$ | $[-0.10, -0.05]$ |
| condition:reward | 0.00 | $[-0.03, +0.02]$ |
| condition:transition | 0.01 | $[-0.02, +0.03]$ |
| reward:transition | $-0.01$ | $[-0.03, +0.01]$ |
| condition:reward:transition | 0.01 | $[-0.01, +0.03]$ |

Table S6: This table reports the group-level coefficients from a hierarchical Bayesian linear regression that seeks to explain pupil diameter at feedback as a function of instruction condition and reward. Instruction condition was coded as $+1$ for story and $-1$ for abstract; reward outcome as $+1$ for reward and $-1$ for no reward; transition as $+1$ for common and $-1$ for rare. CI = credible interval.

The pupil results cannot be explained by different reward frequencies between the two conditions. First, the model included a regressor for reward outcome. Second, performance was similar for both conditions: the mean reward frequency was 0.53 (SD 0.07) for the story compared to 0.54 (SD 0.06) for the abstract condition. Similar performance in the two groups is expected despite differences in strategies because model-free and model-based agents earn similar payoffs in the two-stage task [10]. Therefore, we argue that the larger pupil diameter after feedback in the abstract condition is consistent with greater effort, uncertainty, and/or exploration in that group relative to the story condition. This interpretation is also consistent with the participants' ratings for effort and understanding as well as the pattern of fMRI results.

#### Switch choices at the first stage are associated with larger pupil diameter in the interval between the first and second stages

In a post-hoc, exploratory analysis, we found that the number of stay or switch choices at the first stage correlated with pupil diameter in the four seconds between the first- and second-stage choices for participants in both instruction conditions. We ran a hierarchical Bayesian linear model for the pupil diameter z-scores in this interval that included an indicator for instruction condition (encoded as $-1$ and $+1$), the number of consecutive times the participant repeated the same first-stage choice, mean-centered and quantified on the logarithmic scale[3], and the interaction of these predictors. This model can be written in brms [11] notation as follows:

$$\text{pupil diameter} \sim (1 + \log(\text{repeated choices})|\text{participant}) + \text{condition} * \log(\text{repeated choices}). \quad (5)$$

The effect of the story condition was 0.01 (95% CI $[-0.03, 0.06]$), the effect of repeated choices was $-0.10$ (95% CI $[-0.12, -0.07]$), and the interaction of the story condition and repeated choices was 0.02 (95% CI $[-0.01, 0.04]$).


[3] This is because the effect of repeated choices was assumed to grow more slowly as the number of repetitions increases, that is, the difference between repeating the same choice once versus twice is expected to be larger than the difference between repeating the same choice 10 times versus 11 times.
